# Emergent lag phase in flux-regulation models of diauxie

**DOI:** 10.1101/2023.02.07.527446

**Authors:** Fiona Bate, Yumechris Amekan, Mitya Pushkin, James P.J. Chong, Martin Bees

## Abstract

Lag phase is observed in bacterial growth during a sudden change in conditions: growth is inhibited whilst cells adapt to the environment. Bi-phasic, or diauxic growth is commonly exhibited by many species. In the presence of two sugars, cells initially grow by consuming the preferred sugar then undergo a lag phase before resuming growth on the second. Biomass increase is characterised by a diauxic growth curve: exponential growth followed by a period of no growth before a second exponential growth. Recent literature lacks a complete dynamic description, artificially modelling lag phase and employing non-physical representations of precursor pools. Here, we focus on glucose-lactose diauxie of *Escherichia coli* formulating a rational mechanistic model based on flux-regulation/proteome partitioning with a finite precursor pool that reveals core mechanisms in a compact form. Unlike earlier systems, the characteristic dynamics emerge as part of the solution, including the lag phase, and results accurately reproduce experiments. We show that for a single strain of *E. coli*, diauxic growth yields mathematically optimised growth rates. However, intriguingly, for two competing strains diauxic growth is not always the best strategy. Our description can be generalised to model multiple different microorganisms and investigate competition between species/strains.

## 1 Introduction

Microbial cells show four phases of growth: lag, log (exponential), stationary and death. Lag phase is observed when microorganisms are subject to a sudden change in conditions, such as the introduction of fresh growth media. During lag phase cells adapt to their new environment, synthesising the cellular components necessary for growth. Diauxic growth, first described by Monod [1, 2], occurs when a microorganism is presented with two sugars that can be metabolised. The microorganism first consumes the preferred sugar until that source is almost completely exhausted, only then switching to consume the second food source [2]. There is a lag phase between the two phases of microbial growth on the different food sources which appears to be the result of a trade-off between rapid adaptation to changing growth conditions and supporting a high (and therefore competitive) growth rate [3]. Diauxic growth can be interpreted as a way to maximise growth on two substrates [4, 5]: the sequential use of substrates rather than the simultaneous consumption being beneficial under a wide range of conditions [3]. However, the exact conditions are unclear for which diauxic growth performs better than other strategies, such as consuming both substrates at the same time, albeit at reduced efficiency; in a competitive environment where the two resources are limited, which strain grows most overall?

The underlying molecular interactions governing the response of a microorganism to a change in conditions are complex, although some important regulatory processes have been identified. For example, *E. coli* produces proteins to metabolise lactose only when lactose is present and glucose (the preferred carbon source) is absent. This is achieved through carbon catabolite repression (CCR) and inducer exclusion. CCR is one of the most significant regulatory processes in many bacteria, accounting for 5 − 10% of all bacterial genes [6]. In *E. coli*, CCR is mediated by the prevention of transcriptional activation of catabolic genes in the presence of glucose via the catabolite activator protein (CAP). CAP senses glucose indirectly through the ‘hunger signal’ molecule cyclic adenosine monophosphate (cAMP). Glucose depletion induces *E. coli* to produce more cAMP which binds to CAP, inducing a conformational change that results in binding to DNA, stimulating transcription of the genes involved in lactose metabolism.

The uptake of glucose inhibiting the ability of lactose permease to transport lactose into the cell is known as inducer exclusion [7]. The uptake of glucose by the phosphotransferase system (PTS) is accompanied by the formation of the de-phosphorylated enzyme EIIA^Glc^, which inactivates lactose permease by binding to it [8].

The cooperative coordination of gene expression levels between these two regulatory mechanisms ensures that the preferred carbon source is used first, then metabolism is reconfigured to use the secondary carbon source.

Guanosine 3’,5’-bispyrophosphate (ppGpp), which down-regulates ribosome production and up-regulates amino acid biosynthesis genes, has been found to have an overarching role in coordination of gene expression during glucose-lactose diauxie [9]. The regulation of ribosome synthesis, via ppGpp, is determined by a balance between demand for and synthesis of amino acids. This amino acid flux has been identified as an important factor in the regulation of bacterial growth rate [10]. cAMP, which is important in the regulation of metabolism as noted above, coordinates the expression of catabolic, biosynthetic and ribosomal proteins, ensuring that proteomic resources are spent on distinct metabolic sectors as required in different growth conditions [11].

The mechanisms responsible for reorganisation of gene expression (resource allocation) in microorganisms are generally believed to be optimised by evolution [12]. The optimum mechanism will depend on the growth environment. For example, in a non-competitive environment the maximisation of growth yield is thought to provide an advantage [12] whereas when there is competition for resources, maximising growth rate will give a competitive advantage [13].

Recent theoretical studies on resource allocation have focussed on maximizing growth rate [10, 14]. Scott et al [10] used a coarse-grained model of the cell to show that maximum growth rate is acheived at a specific value of the ribosomal protein fraction through maximisation of the amino acid flux. The amount of protein in the cell was assumed constant and divided into related sectors (proteome partitioning): ribosomal proteins and metabolic proteins. Increasing the number of ribosomes therefore decreases metabolic enzyme levels. Their optimisation control strategy was based on the amino acid pool size, assumed to be signalled via ppGpp, controlling the fraction of total protein synthesis producing ribosomes [10]. Similar models of resource allocation optimisation include energy constraints in addition to constraints on the proteome [15, 16].

The above studies involve steady state models, describing an environment that is stable over a long period of time. However, on the whole a microorganism is subject to a fluctuating range of growth conditions in it’s natural environment. This has motivated the formulation of dynamic resource allocation models [5, 12, 17, 18, 19, 20]. Kremling et al [20] present an ensemble of different models all showing diauxic behaviour. By qualitatively comparing model predictions they offer an insight into the variety of mechanisms that have been proposed to play a role in CCR. Basan et al [19] invesitgated shifts between two single carbon sources reporting that long lag phases are due to the depletion of key metabolites and resulting metabolic bottlenecks. Pre-shift growth rates were varied by using different carbon sources and their model of sequential flux limitation predicts a linear relationship between lag time and pre-shift growth rate. A stochastic simulation model presented by Chu and Barnes [3] shows that it is impossible to shorten the lag phase without reducing the long term growth potential. Premature activation of the secondary metabolism shortens the lag but causes costs to the cell thus reducing the growth rate on the preferred substrate. They predict, using simulated evolution, that the lag phase will evolve to be longer in environments where switching is less likely and shorter in frequently changing environments. Erickson et al [18] present a kinetic flux-controlled regulation model that quantitatively describes adaptation dynamics based on the dynamic reallocation of proteomic resources. The time evolution of gene expression is determined by regulation functions whose form is derived from steady-state growth laws. There are limitations on the validity of these regulation functions and in addition the model predicts constant proportionality between growth rate and substrate uptake rate, which is not observed experimentally during log-phase growth.

In this study we extend and modify the model of Erickson et al [18] to include accurate prediction of biomass growth and substrate uptake during an initial lag-phase and during diauxic shift. We develop a coarse-grained model which uses qualitative knowledge of the molecular processes and a flux balance approach. We have avoided the excessive complication of other models [5] explicitly so that we do not have large numbers of unmeasurable parameters. Unknown kinetic parameters in the model description are related to measurable kinetic parameters to minimise the need for fitting. Unlike many mathematical models describing lag-phase [21], we do not introduce an artificial lag parameter to control the length of the lag. Instead the duration of the lag-phase is determined by the initial structure of the microorganism’s proteome.

We present a rational description, based on experimentally measurable parameters, which reproduces all principal features of the growth curve of *E. coli* during glucose-lactose diauxie. Both the lag phase and log phase of bacterial growth emerge as part of the solution. Such a description (summarised in Section 2.3) can be used to demonstrate the relative merit of diauxic growth over the whole growth period and explore other growth strategies.

## 2 Flux-controlled regulation of anabolism and catabolism

To model flux-controlled regulation (FCR) we shall adopt the modelling formalism of Erickson et al [18], develop a rational mathematical approach to address modelling inconsistencies and extend the description to describe physical aspects of precursor and amino acid pools.

### 2.1 Original FCR model

The FCR model due to Erickson et al [18] describes the time evolution of gene expression and biomass growth during carbon upshifts and downshifts. The model balances carbon influx and protein synthesis flux via changes to the average translation rate, *σ*, which is set by the size of a pool of central precursors including ketoacids and amino acids. Which proteins are produced (catabolic enzymes/ribosomes) is determined by regulation functions whose form is derived from steady-state growth laws. The central assumption of this model is that the time-dependence of the regulation functions during growth transitions depends solely on changes to the translation rate.

#### 2.1.1 Limitations of the original FCR model

The regulation functions defined in [18] are undefined for a particular value of the translation rate, which we will call *σ*_*P*_, and for *σ > σ*_*P*_ the regulation functions incorrectly are negative. Although values of *σ* ≥ *σ*_*P*_ do not occur during steady-state growth they can occur during growth transitions. To remove this inconsistency and provide a firmer theoretical foundation we derive our regulation functions directly, associated with a mathematical optimization of the growth rate (see Section 2.2.5).

The original FCR model [18] states that, on the time scale of interest, all fluxes are balanced. This balance is achieved by assuming that the translation rate adjusts abruptly with any changes to carbon influx (due to changes in substrate availability or the concentration of a key enzyme). However, for small values of the ribosome mass fraction or large carbon influx this can lead to large, physically unrealistic translation rates. We reason that as the translation rate depends on the size of the precursor pool, which is finite, the rate must be limited. Therefore, we shall include this limitation in our model (see Section 2.2.3).

Moreover, requiring flux balance in the above way results in the protein synthesis rate, and hence biomass growth rate, only depending on the catabolic protein mass fraction: the ribosome mass fraction drops out of the equations. The resulting constant proportionality between growth rate and substrate uptake (the constant biomass yield) predicted by the model of Erickson et al [18] does not agree with experimental observations. Our data, which we present in Section 3.1, shows that during an initial lag phase the ratio of growth rate to substrate uptake rate is significantly less than it is during the subsequent log-phase growth: the biomass yield is not constant. This suggests that growth is not being limited solely by the catabolic proteins, as this would also limit substrate uptake, but must depend on the levels of other key proteins.

#### 2.1.2 Factors limiting growth during the initial lag phase

Prior to the diauxie experiment (a full description of which is given in Section 3.1) *E. coli* was grown on Luria-Bertani broth (LB) which contains carbon sources and amino acids essential for growth. Cells of *E. coli* growing in LB can import amino acids directly and therefore do not need to use anabolic proteins to build amino acids. Indeed, it has been found experimentally that *E. coli* grown in LB show much lower levels of many genes involved in the amino acid biosynthetic pathways than those grown in minimal media [22]. Therefore, we propose that the lag phase occurring when *E. coli* switches from growth on rich LB to minimal media is caused by a lack of the anabolic proteins needed for the biosynthesis of amino acids. To investigate this we extend the original FCR model to include an amino acid synthesis flux.

### 2.2 Modified and extended FCR model

External substrates, *S*_*j*_, are consumed by a microorganism, *X*. Inside the microbial cell catabolic enzymes break the substrate down into precursors. Anabolic proteins combine precursors to form amino acids that are subsequently incorporated by ribosomes into proteins required for growth. The relative amounts of the different enzymes and proteins required are determined by the growth conditions and substrates being consumed.

We construct a mathematical description of this process incorporating proteome partitioning, flux-controlled regulation and allocation of protein synthesis via optimisation of the growth rate.

#### 2.2.1 Proteome partitioning

Using an established model of proteome partitioning [14, 23] we split the total protein content of the cell into different sectors, each composed of proteins whose expression levels show similar growth rate dependency in different growth conditions. The growth rate dependent sectors of the proteome are ribosome-affiliated proteins, *R*, enzymes relating to carbon import and metabolism, *C*, anabolic enzymes related to the production of amino acids, *A*, and an ‘uninduced’ sector, *U*, which generally decreases with decreasing growth rate [11]. The rest of the proteome, *Q*, is growth rate independent and its mass fraction is non-zero and constant. It follows that

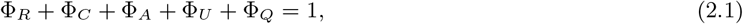

where Φ_*i*_ is the mass fraction of sector *i*. The minimum mass fraction of each sector, Φ_*i*,0_, is assumed to be growth rate independent [11] so that for each sector the growth rate dependent part is given by *ϕ*_*i*_ = Φ_*i*_ − Φ_*i*,0_. Thus, in terms of the growth rate dependent parts of the mass fractions, equation (2.1) becomes

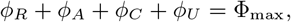

where Φ_max_ = 1 − Φ_*Q*_ − Φ_*R*,0_ − Φ_*A*,0_ − Φ_*C*,0_ − Φ_*U*,0_ < 1 is a constant. This can be further simplified by noting that the uninduced sector of the proteome is found to be related to the ribosomal sector (under *C* and *A* limitation) such that *ϕ*_*U*_ = *εϕ*_*R*_ (You et al [11] establish *ε* = 0.3). It follows that

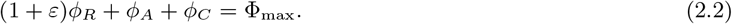

During the log phase of growth of bacterial cells, the rate of cell proliferation (the growth rate) and the expression levels of key proteins are linearly correlated [11, 14, 18]. Each protein sector is assumed to be regulated as a whole [11, 24] so *ϕ*_*i*_ is proportional to the expression level of a key protein in sector *i*, and thus to the growth rate.

Denoting the value of the mass fraction during the log phase by 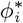 it follows that

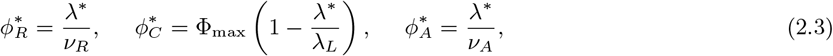

where *λ*^*^ is the constant growth rate of the *E. coli* cells in log phase and *ν*_*R*_, *λ*_*L*_ and *ν*_*A*_ are constants (see Appendix A for further details).

#### 2.2.2 Flux-controlled regulation

The core mechanisms represented in our model are shown in Figure 1. The microorganism takes up substrates and breaks them down into carbon precursors. These precursors, together with other essential nutrients, combine to supply the cell with a pool of amino acids. The amino acids are utilised by ribosomes to produce proteins, *Z*. The rate of protein synthesis depends on the concentration of ribosomes, *R*, and the average translation rate, *σ*_*A*_, so that d*Z/*d*t* = *σ*_*A*_*R*. The total mass of protein as a fraction of total biomass is relatively constant [18]. Therefore, the total biomass concentration, *X*, is related to the total protein concentration by *Z* = *pX*, where the constant *p* is the fraction of biomass that is protein. It follows that

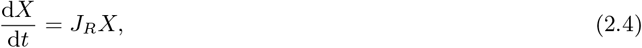

where *J*_*R*_ = *σ*_*A*_Φ_*R*_ represents the protein synthesis flux and Φ_*R*_ = *R/*(*pX*). Analogous to *J*_*R*_ the amino acid synthesis flux is given by 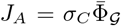, where *σ*_*C*_ is the average amino acid synthesis rate and 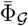 is the rescaled mass fraction of a key anabolic protein, 𝒢. (We rescale Φ _𝒢_ with a factor proportional to Φ_max_ to remove an unknown constant from the equations, details are given in Appendix A.3). The relationship between 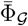 and the mass fraction of the total amino acid sector, Φ_*A*_, where Φ_*A*_ = *A/*(*pX*), is discussed in Section A.3 of the Appendix, with equation (A.4) giving the explicit dependence.

**Figure 1:**
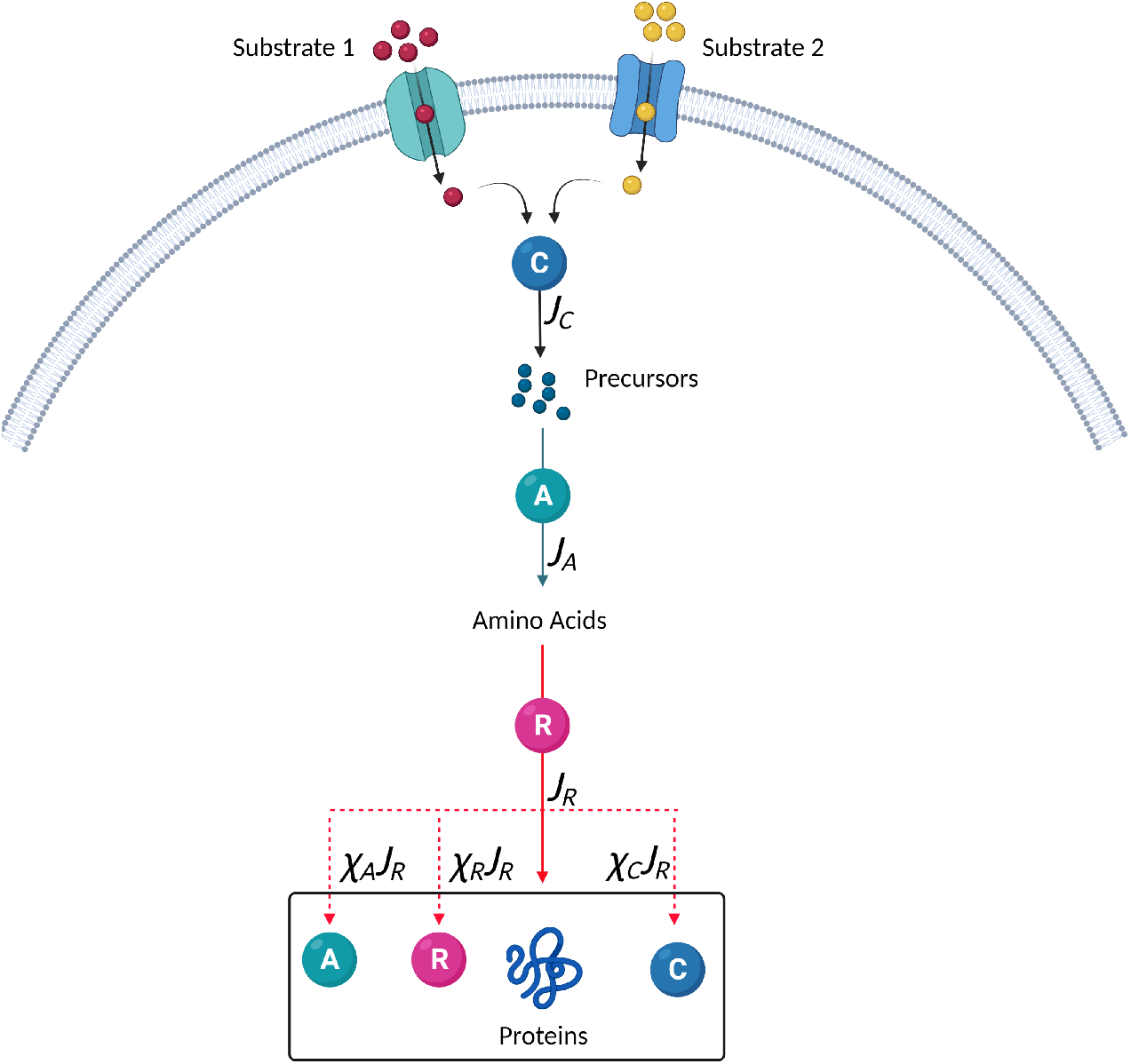
Flux-controlled regulation model. External substrates, *S*_*j*_, are taken in and then broken down by catabolic enzymes, the C-sector, to supply a pool of carbon precursors. Changes in the concentration of the substrates and enzyme result in changes to the carbon influx, *J*_*C*_. Other essential nutrients, including nitrogen, combine with these carbon precursors and are built up by anabolic proteins, the A-sector, to form amino acids. The flux of amino acid synthesis is given by *J*_*A*_. A balance between *J*_*A*_ and *J*_*C*_ is achieved through changes to the average amino acid synthesis rate, *σ*_*C*_, which in turn depends on the size of the precursor pool. The amino acids are “consumed” by ribosomes, the R-sector, in protein synthesis. The flux of protein synthesis is given by *J*_*R*_. A balance between *J*_*R*_ and *J*_*A*_ is achieved through changes to the average translational activity, *σ*_*A*_, which depends on the size of the amino acid pool. The regulation functions *χ*_*R*_, *χ*_*A*_ and *χ*_*C*_ determine the amount of ribosomal, anabolic and catabolic proteins, respectively, that are produced. Allocation of protein synthesis is regulated, via ppGpp and cAMP [9, 10, 11], in response to changes to the precursor and amino acid pools. Under given growth conditions, there is an optimum level for each protein that will maximise the growth rate. During growth transitions the proteins are not at optimum levels, leading to changes in the precursor and amino acid pools and a non-optimum growth rate. In the model the regulation functions are obtained by mathematically optimising the growth rate. (Image created with BioRender.com)

The carbon influx rate, *J*_*C*_, is proportional to the rate of substrate uptake. As the C-sector as a whole is shared between all substrates we base the substrate uptake equation on Michaelis-Menten kinetics for multiple substrates (see Appendix B for details). For the case where there are *N* substrates, with concentrations {*S*_*j*_} = {*S*_1_, *S*_2_, …*S*_*N*_},we have

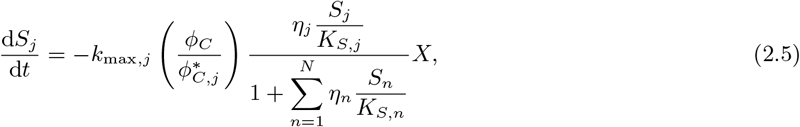

where, as before, *ϕ*_*C*_ = Φ_*C*_ − Φ_*C*,0_ and Φ_*C*_ = *C/*(*pX*). The constants *k*_max,*j*_ and *K*_*S,j*_ are the maximum uptake rate and the affinity constant for substrate *j* respectively. The function *η*_*j*_ ({*S*_*j*_}), where 0 ≤ *η*_*j*_ ≤ 1, depends on which substrates are present in the system and determines whether a specific substrate is being consumed. For example, when modelling glucose-lactose diauxie, glucose will always be consumed so we set *η*_gl_ = 1. However, lactose is only consumed when the concentration of glucose drops sufficiently. The point at which lactose uptake switches on is not very well defined but we require *η*_la_ → 0 when the glucose concentration, *S*_gl_, is large and *η*_la_ → 1 as *S*_gl_ → 0. This can be modelled by setting

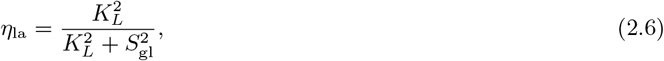

where *K*_*L*_ is a constant. This choice of function facilitates a smooth transition between repressing lactose uptake when glucose concentrations are high and no repression of lactose uptake at zero glucose concentration. Glucose levels must drop so that *S*_gl_ ≈ *K*_*L*_ before lactose begins to be consumed.

We define *Y*_*C,j*_ to be the yield of carbon precursors from *S*_*j*_, so, obtaining the substrate uptake rate from equation (2.5), it follows that the carbon influx rate due to substrate *S*_*j*_ is given by

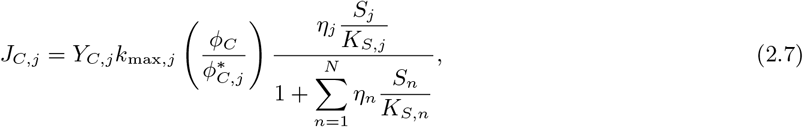

with the total carbon flux 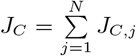. Note that we do not assume that *Y*_*C,j*_ is constant, as is the case in [18], as this would result in the biomass yield, *Y*_*j*_, being constant, which is inconsistent with experimental observations (as discussed in Section 2.1.1). In our model *Y*_*C,j*_ and, therefore, the biomass yield, *Y*_*j*_, depend on the growth conditions and proteome structure as we now show.

#### 2.2.3 The finite precursor pool size

When growth conditions change, the amount of carbon available to enter the growth pathway (shown in Figure 1), via the carbon influx, *J*_*C*_, is affected. An abrupt upshift in substrate quality could cause *J*_*C*_ to increase suddenly, resulting in a sudden increase in the production rate of carbon precursors. The level of the A-sector proteins cannot increase abruptly (as protein synthesis rates are proportional to the growth rate) and a bottleneck will occur in the growth pathway. This could be dealt with by abruptly increasing the amino acid synthesis rate, *σ*_*C*_, as in [18], but accounting for large changes in *J*_*C*_ in this way requires setting unrealistically high values for *σ*_*C*_. Instead we note that the size of the precursor pool is limited by a cell’s capacity, there being only finite space within a cell. Thus the abundance of carbon precursors is limited which, as *σ*_*C*_ depends on the abundance of carbon precursors, in turn limits the value of *σ*_*C*_. (Similarly, the translational activity, *σ*_*A*_, will have a maximum value.) To maintain flux balance we propose that the carbon entering the growth pathway, *J*_*C*_, is limited. This is achieved by allowing the yield of carbon precursors, *Y*_*C,j*_, to change as growth conditions change. Note that the substrate that is broken down but does not enter the growth pathway will be released as product (which we do not explicitly model). This is the case whether *Y*_*C,j*_ is constant, as in [18], or changing, as in this model.

We let *P*_*C,j*_ represent the concentration of carbon precursors added to the precursor pool by the flux *J*_*C,j*_, defined in equation (2.7), and *P*_*A,j*_ represent the amino acids subsequently synthesised from *P*_*C,j*_. The combined size of the carbon precursor and amino acid pools can therefore be written as

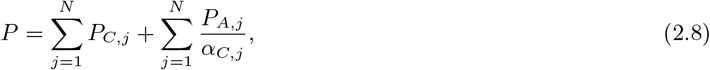

where *α*_*C,j*_ is a constant conversion factor from *P*_*C,j*_ to *P*_*A,j*_. There is a maximum value of *P* that can be maintained in the cell and we denote this by *K*. This constant, *K*, is analogous to the carrying capacity in population dynamics, the maximum population size that can be sustained in a given growth environment. In population dynamics the growth rate is limited by the carrying capacity, with growth tending to zero as the population size tends towards the carrying capacity. Here we limit the fluxes entering the carbon precursor pool so that *J*_*C,j*_ → 0 as *P* → *K*. We have

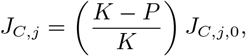

where *J*_*C,j*,0_ is the carbon flux from substrate *j* when *P* = 0 given by

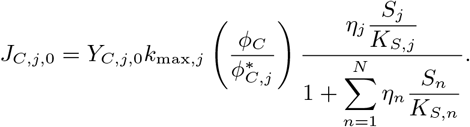

The constant *Y*_*C,j*,0_ is the yield of carbon precursors from *S*_*j*_ when *P* = 0. To simplify notation we introduce the function

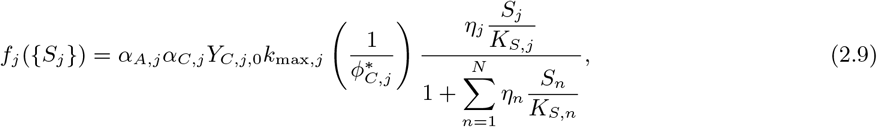

so that

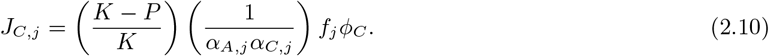

To keep the number of variables in the model to a minimum we want the carbon influxes *J*_*C,j*_ to be defined only in terms of the substrate concentrations and protein mass fractions. This means we need to know *P*, and therefore *P*_*C,j*_ and *P*_*A,j*_, only in terms of the substrate concentrations and protein mass fractions. This is done by considering flux balance.

The amino acid synthesis flux is given by 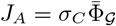, as discussed in section 2.2.2, where *σ*_*C*_ = *σ*_*C*_ ({*P*_*C,j*_}) depends on the abundance of carbon precursors. For simplicity, we take a linear dependence, setting *σ*_*C*_ = ∑_*j*_ *α*_*C,j*_ *k*_*C,j*_ *P*_*C,j*_, where *k*_*C,j*_, the uptake rate of *P*_*C,j*_, is a constant. The amino acid synthesis flux due to substrate *j* is therefore given by 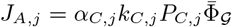. Similarly, as the total protein synthesis flux *J*_*R*_ = *σ*_*A*_Φ_*R*_, we take *σ*_*A*_ = ∑_*j*_ *α*_*A,j*_ *k*_*A,j*_ *P*_*A,j*_, where the constant *k*_*A,j*_ is the uptake rate of *P*_*A,j*_ and *α*_*A,j*_ is a constant conversion factor from *P*_*A,j*_ to protein, and obtain the protein synthesis flux due to substrate *j* as *J*_*R,j*_ = *α*_*A,j*_ *k*_*A,j*_ *P*_*A,j*_ Φ_*R*_.

The rates of change of *P*_*C,j*_ and *P*_*A,j*_ in terms of the fluxes, *J*_*C,j*_, *J*_*A,j*_ and *J*_*R,j*_ are given by

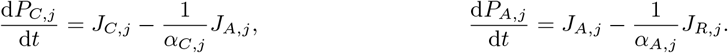

To achieve flux balance, changes in *P*_*C,j*_ and *P*_*A,j*_ are assumed to take place over a relatively fast time scale, so that d*P*_*C,j*_ */*d*t* = d*P*_*A,j*_ */*d*t* = 0. Essentially this means that on the timescale of interest all fluxes balance so that

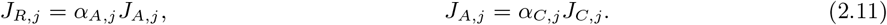

Substituting for *J*_*R,j*_, *J*_*A,j*_ and *J*_*C,j*_ in equations (2.11) we obtain a system of 2*N* equations in terms of *P*_*C,j*_ and *P*_*A,j*_. These can be solved to give *P*_*C,j*_ and *P*_*A,j*_ in terms of the substrate concentrations and protein mass fractions. From these we can then work out *P*, using equation (2.8), and substituting for *P* into (2.10) we obtain

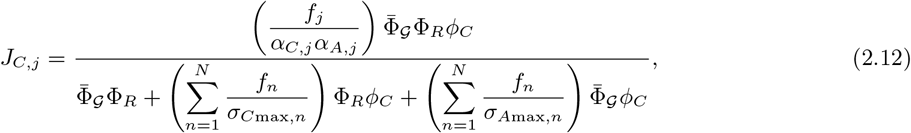

where *σ*_*A*max,*n*_ = *α*_*A,n*_*α*_*C,n*_*k*_*A,n*_*K* and *σ*_*C*max,*n*_ = *α*_*A,n*_*α*_*C,n*_*k*_*C,n*_*K* are, respectively, the maximum translation rate and maximum amino acid synthesis rate when only substrate *n* is being consumed. Full details of the derivation of equation (2.12) are given in Appendix C.

Comparing equation (2.12) with the definition of *J*_*C,j*_ given by equation (2.7), it follows that

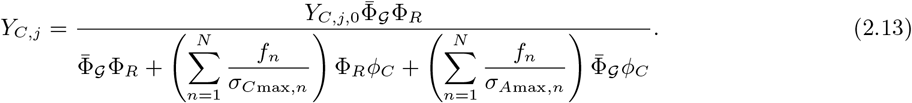

This equation describes how the yield of carbon precursors changes with the substrate concentrations (through *f*_*j*_) and protein mass fractions.

We now use the expression we have derived for *J*_*C,j*_, given by equation (2.12), and the flux balance equations (2.11) to derive an equation for biomass growth.

#### 2.2.4 Biomass growth

The equation for biomass growth in terms of the protein synthesis flux, 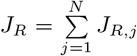, is given by equation (2.4). From flux balance we have *J*_*R,j*_ = *α*_*A,j*_ *α*_*C,j*_ *J*_*C,j*_, with *J*_*C,j*_ given by equation (2.12). It follows that

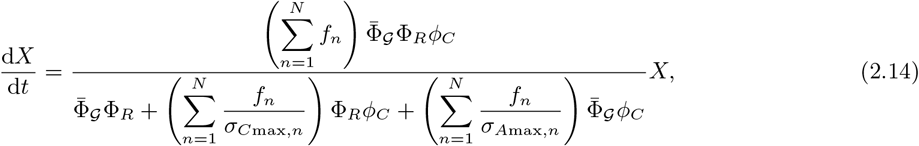

with *f*_*j*_, given by equation (2.9). Note that the growth rate

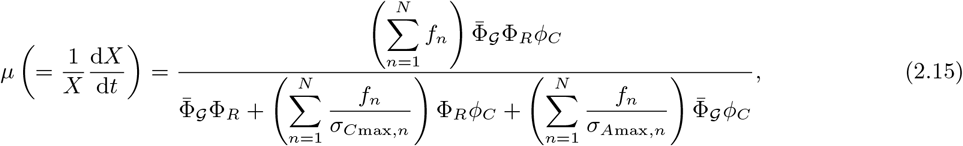

is not directly proportional to the substrate uptake rate and depends on the mass fractions of each of the growth dependent proteome sectors. For small Φ_*R*_ the growth rate is limited by Φ_*R*_ and similarly for 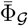 and *ϕ*_*C*_. Crucially, for fixed substrate concentration (constant *f*_*j*_), the growth rate 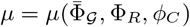 has a unique maximum value at specific values of 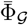, Φ_*R*_ and *ϕ*_*C*_. We hypothesise that during the log-phase cells grow at this optimal rate. This hypothesis uniquely defines the values of the unknown constants in equations (2.14) and (2.15), *σ*_*C*max,*j*_, *σ*_*A*max,*j*_ and *α*_*A,j*_ *α*_*C,j*_ *Y*_*C,j*,0_ (this latter combination of constants appears in the definition of *f*_*j*_). In terms of experimentally measurable parameters we find

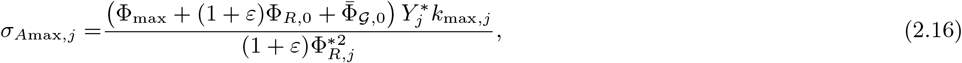

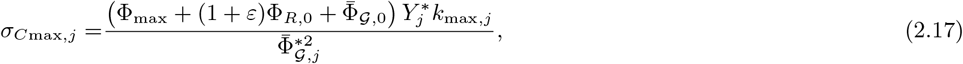

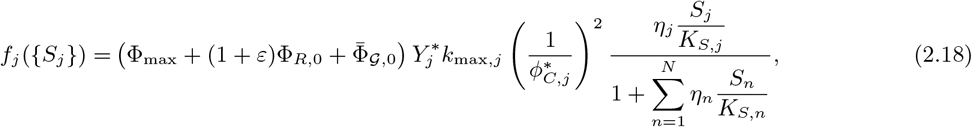

where the constant 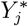 is the measured biomass yield during log-phase growth on substrate *j*. Full details are given in Appendix D.

#### 2.2.5 Allocation of the protein synthesis flux

Allocation of protein synthesis is regulated, via ppGpp and cAMP, in response to changes in the precursor and amino acid pools [9, 10, 11]: protein levels are adjusted until pool sizes are optimal. Variations in pool sizes manifest as changes to the growth rate (the size of precursor and amino acid pools being explicitly included in the derivation of the growth rate equation (2.15), see Section 2.2.3) and it follows that when precursor and amino acid pool sizes are optimal the growth rate is maximal. We therefore allocate the protein synthesis flux through regulation functions that adjust the level of proteins until the growth rate is maximal.

Proteins belonging to the growth independent sector of the proteome, *Q*, are produced as a constant fraction, Φ_*Q*_, of total protein production. Proteins belonging to the growth dependent proteome sectors are produced in varying amounts depending on the current state of the proteome and the growth conditions. The *R, A* and *C* sectors of the proteome are each composed of a growth independent part and a growth dependent part. The growth independent parts are produced at a constant fraction of total protein produced as for sector *Q*. For the growth dependent parts we define regulation functions *χ*_*R*_, *χ*_*C*_ and *χ*_*A*_ to represent the fraction of the total amount of protein produced that is R-sector, C-sector, and A-sector protein respectively so that

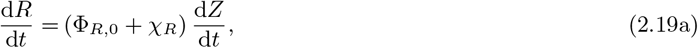

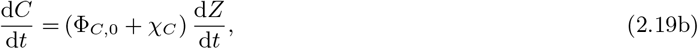

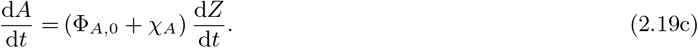

As the total amount of protein is fixed the regulation functions are constrained by

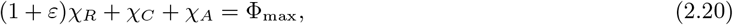

and we also require *χ*_*R*_ ≥ 0, *χ*_*C*_ ≥ 0 and *χ*_*A*_ ≥ 0 as protein is not recycled or destroyed (this is a simplifying assumption of our model). Using the relationships between biomass concentration and total protein concentration, and protein concentrations and protein mass fractions, given in section 2.2.2, we rewrite equations (2.19) in terms of the growth dependent protein mass fractions (details given in Appendix E). We have

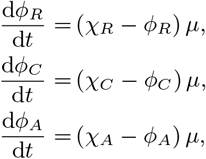

with the growth rate, *μ*, given by equation (2.15). From here on we will not solve explicitly for *ϕ*_*A*_ as its value is determined from *ϕ*_*R*_ and *ϕ*_*C*_ using equation (2.2).

When external substrate concentrations are constant the substrate dependent functions *f*_*j*_ have constant values and the mass fractions are assumed to take values that maximise *μ* for the given conditions. When the external substrate concentrations change, the mass fractions adjust, via the regulation functions, so that for the new values of *f*_*j*_ the growth rate is maximised. Using the standard calculus result that the greatest rate of increase of a function at a given point is in the direction given by the gradient of that function at that point, we note that the optimum way to get to the maximum value of *μ* is to set

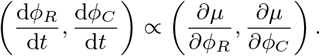

To ensure that the regulation functions are always positive we define

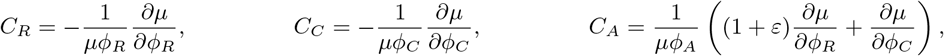

and set

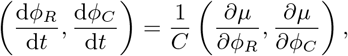

where *C* = max{1, *C*_*R*_, *C*_*C*_, *C*_*A*_}. (Note that if *ϕ*_*R*_ = 0, *C*_*R*_ is undefined and we must set *C*_*R*_ = 1 before working out *C*, similarly for *ϕ*_*C*_ and *ϕ*_*A*_.) The resulting regulation functions

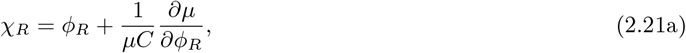

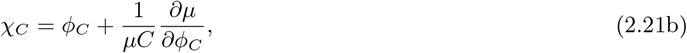

are a mathematical representation of protein synthesis regulation via ppGpp and cAMP: protein levels are adjusted to optimise growth rate. The regulation functions are never negative and the constraint on protein production, given by equation (2.20), is always satisfied.

### 2.3 Governing equations

In summary, we have constructed a mechanistic model describing the time evolution of biomass growth, substrate concentration and gene expression during carbon upshifts and downshifts. Phases of microorganism growth emerge from the dynamics of the proteome, rather than being switched on/off at a particular time. The model incorporates proteome partitioning, flux-controlled regulation and optimal allocation of protein synthesis. The governing equations are

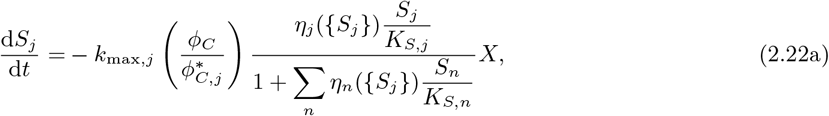

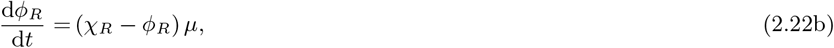

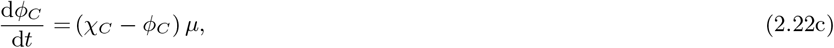

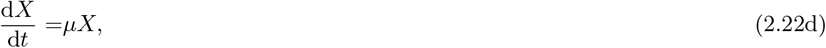

with the growth rate, *μ* = *μ*({*S*_*j*_}, *ϕ*_*R*_, *ϕ*_*C*_), given by equation (2.15). An overview of the model variables and parameters is given in Tables 1 and 2.

**Table 1:**
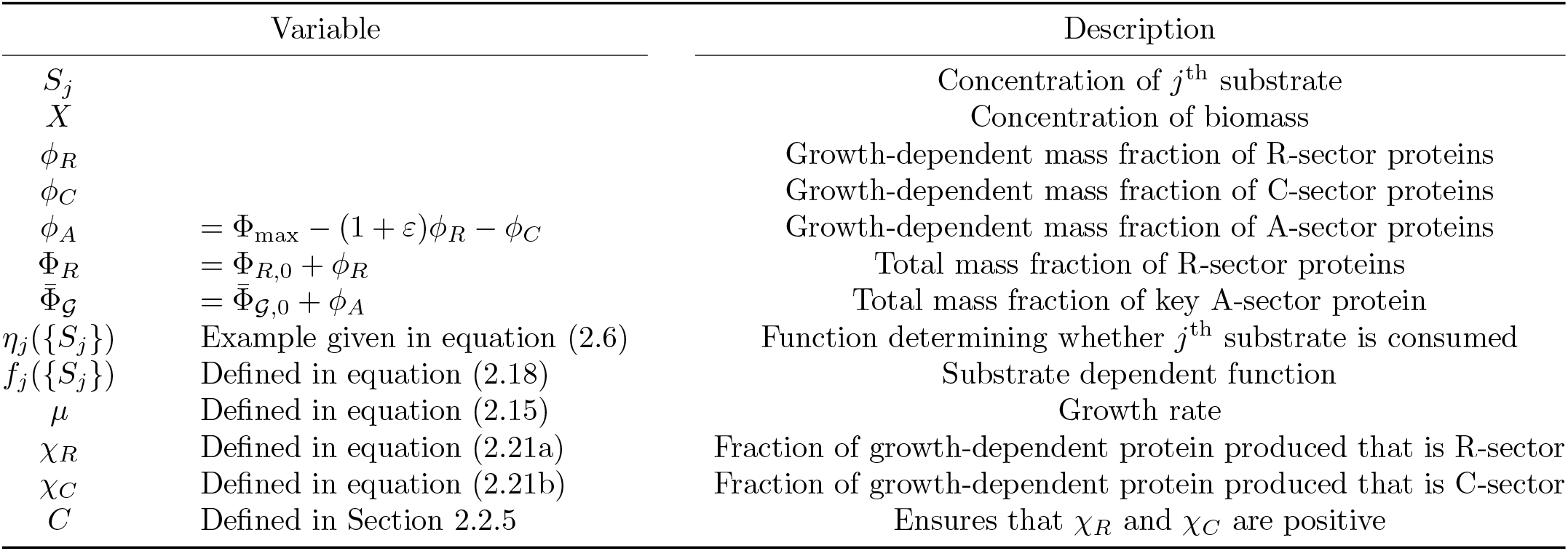
Overview of model variables. A full description and derivation of variables is given in the text.

**Table 2:** Overview of model parameters. A full description and derivation of parameters is given in the text.

| Parameter | Description |
| --- | --- |
| $Y_j^*$ | Biomass yield measured during log-phase growth on substrate $S_j$ |
| $k_{\max,j}$ | Maximum uptake rate of $S_j$ |
| $K_{S,j}$ | Half saturation constant on $S_j$ |
| $\nu_R$ | Fitted parameter in growth law |
| $\lambda_L$ | Fitted parameter in growth law |
| $\nu_A = \left( \frac{\Phi_{\max}}{\lambda_L} - \frac{(1+\varepsilon)}{\nu_R} \right)^{-1}$ | Fitted parameter in growth law |
| $\Phi_{\max}$ | Total combined mass fraction of growth dependent proteins |
| $\varepsilon$ | Constant relating uninduced sector to R-sector |
| $\phi_{R,j}^* = \frac{Y_j^* k_{\max,j}}{\nu_R}$ | Value of $\phi_R$ during log-phase growth on substrate $S_j$ |
| $\phi_{C,j}^* = \Phi_{\max} \left( 1 - \frac{Y_j^* k_{\max,j}}{\lambda_L} \right)$ | Value of $\phi_C$ during log-phase growth on substrate $S_j$ |
| $\phi_{A,j}^* = \frac{Y_j^* k_{\max,j}}{\nu_A}$ | Value of $\phi_A$ during log-phase growth on substrate $S_j$ |
| $\Phi_{\mathcal{R},0}$ | Minimum ribosomal mass fraction |
| $\Phi_{\mathcal{G},0}$ | Minimum mass fraction of key A-sector protein |
| $\sigma_{A\max,j} = \frac{(\Phi_{\max} + (1+\varepsilon)\Phi_{R,0} + \bar{\Phi}_{\mathcal{G},0}) Y_j^* k_{\max,j}}{(1+\varepsilon)\Phi_{R,j}^{*2}}$ | Maximum translation rate when consuming $S_j$ |
| $\sigma_{C\max,j} = \frac{(\Phi_{\max} + (1+\varepsilon)\Phi_{R,0} + \bar{\Phi}_{\mathcal{G},0}) Y_j^* k_{\max,j}}{\bar{\Phi}_{\mathcal{G},j}^{*2}}$ | Maximum amino acid synthesis rate when consuming $S_j$ |

The exact mechanism underlying the inhibition of substrate uptake is not made explicit in the model, making it flexible and applicable to many processes. In addition, the description can be generalised to model multiple different microorganisms, facilitating investigation of competition between different species or strains.

We now apply this model to the particular case of glucose-lactose diauxie of *E. coli*, comparing numerical results to preliminary experimental data.

## 3 Results

In order to parameterize and test the mathematical model we reproduced the classic *E. coli* glucose-lactose diauxie experiment.

### 3.1 Methods

We recreated the glucose-lactose diauxie experiment of Mostovenko et al [25], using mixed *E. coli* strains. The following two strains were employed: *E. coli* MV1717 (MG1655 *lac*^*+*^ containing chromosome-encoded, inducible CDI-msfGfp, chloramphenicol (Cm) resistance) and *E. coli* MV1300 (MG1655 delta lacZYA; kanamycin (Kan) resistance). Strain MV1717 can grow on lactose (*lac*^*+*^), while MV1300 cannot utilise lactose (*lac*^*-*^) as it is missing the lacY gene that encodes lactose permease, a membrane transporter that pumps lactose into cells. This characteristic was confirmed by growth on MacConkey agar, as shown in Figure 2, where MV1717 (*lac*^*+*^) colonies grow pink and MV1300 (*lac*^*-*^) colonies grow colourless (white).

**Figure 2:**
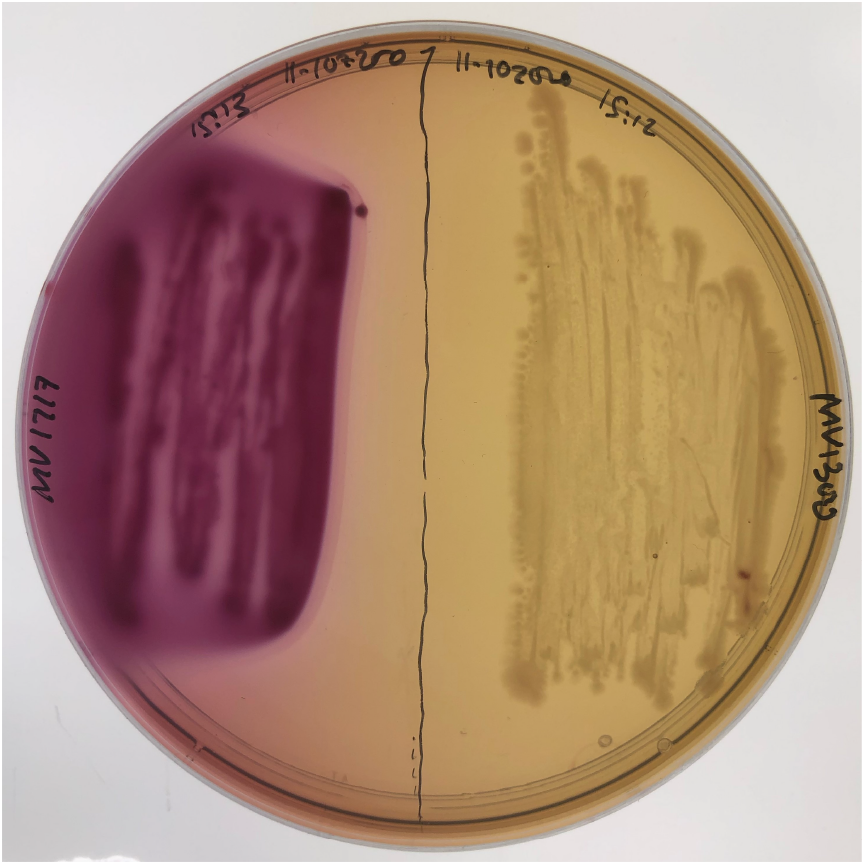
The two utilised *E. coli* strains growing on differential MacConkey agar. Lactose fermenters grow red or pink, cells unable to utilise lactose do not change colour. Left: MV1717 a lactose fermenting (*lac*^*+*^) strain. Right: MV1300 a non-lactose utilising (*lac*^*-*^) strain

Strains were grown separately overnight on Luria-Bertani agar (LB-agar Miller; LMM0204, Formedium, Hunstanton, UK) at 37°C. A single colony of each strain was then grown overnight in 50 mL Falcon tubes containing 25 mL LB medium (LB broth, Miller; BP9723-500, Fisher BioReagents, Loughborough, UK) at 37°C with 120 rpm orbital shaking. Both the LB-agar and LB broth contained antibiotics: 30 μg/mL Kan for MV1300 and 34 μg/mL Cm for MV1717 respectively. The use of antibiotics was necessary for the qPCR we carried out which targeted plasmids carried by these strains that were used for quantification.

Biomass was measured using an established OD vs cell density relationship for *E. coli* [26]. (Note that whilst OD will likely depend on the geometry of cells, granularity and other aspects, it is commonly assumed to be proportional to biomass.) When the optical density at 600 nm (OD600) reached 1.4 (∼ 1.1 *×* 10^9^ cells/mL), the biomass from each tube was harvested via centrifugation (Centrifuge 5810 R, Eppendorf) at 1940*×* g, 37° C. Supernatants were removed and the pellet was resuspended in 10 mL of warm (37°C) filter sterile 1x phosphate buffered saline (PBS 20-7400-10, Severn Biotech Ltd.). The biomass was spun again with the same parameters. The PBS was removed and the pellet was resuspended in 10 mL of 1x Morpholinepropanesulfonic acid (MOPS) minimal medium (Teknova, Hollister, CA, USA).

The strains were mixed in 1:1 ratio (v/v) prior to inoculation and OD600 measured (the inoculum OD600 was 6.82). The mixed culture was used to inoculate a 3 L glass autoclavable bioreactor (Applikon Biotechnology, Delft, The Netherlands) with 1 L of 1x MOPS minimal medium (Teknova, Hollister, CA, USA) containing 0.5 g/L glucose and 1.5 g/L lactose as the only carbon sources [9, 25]. Bioreactor temperature was maintained at 37°C (*±*0.3°C) via a recirculating water bath (OLS200, Grant Instruments). Culture pH was monitored and logged via a Bio Controller (ADI 1010, Applikon Biotechnology, Delft, The Netherlands) and maintained at pH 7.2 *±* 0.05 by addition of 2 M NaOH. Dissolved oxygen was maintained above 20% saturation by adjusting agitation speed in the range of 270 – 500 rpm (Motor Controller, ADI 110, Applikon Biotechnology, Delft, The Netherlands) with fixed 1 L/min air flow [9].

To monitor cell density and glucose and lactose concentration, 2 mL samples were collected every 30 minutes before and after diauxie and every 10 minutes near and during the diauxic shift, as described in [25]. *E. coli* growth was measured by assessing OD600 using a Thermo Spectronic Biomate 3 UV-Visible spectrophotometer (ThermoFisher Scientific, UK) zeroed against an uninoculated growth medium blank. For large values of OD600 (> 0.4), we calculated OD600 based on samples that were diluted in media and remeasured. The concentrations of glucose and lactose were assayed using enzymatic kits (CBA086, Sigma-Aldrich and K624, BioVision, respectively). Aliquots of cells were also cultured on MacConkey agar and incubated at 37°C overnight for differentiation and enumeration of lactose and non-lactose fermenting strains.

Cell density and glucose and lactose concentration measurements allowed the accurate establishment of the initial lag phase (caused by the switch from growth on rich LB to minimal media) and the onset of diauxic growth, see Figure 3. During the initial lag phase (up to around 200 minutes) substrate is taken up (the glucose level declines; see Figure 3(b)) but the growth rate is significantly less (as is evident from a log-linear plot, not shown). The longer initial lag phase observed in experiments 1 and 2 is likely due to the fact that the two cultures used to inoculate experiments 1 and 2 were slightly older (20h post inoculation) compared to experiment 3 (18h post inoculation). Diauxie began when the culture reached OD600 of ∼ 0.5 or a density of approximately 4×10^8^ cells/mL [26] and was indicated by a 20 − 30 minute plateau in the growth curve (Figure 3). The OD600 of the diauxic shift was comparable in three experiments (OD600 of 0.52, 0.59, 0.55), see Figure 3(a) and [25]. The onset of diauxie corresponded to exhaustion of glucose in the growth media. Lactose was depleted at around 250 minutes after the diauxic shift and growth reached stationary phase when OD600 ∼ 2. Data for qPCR indicate that MV1717 and MV1300 are present in approximately equal numbers while glucose is available in the media but that only MV1717 continues to grow exponentially after the lag phase associated with the switch to fermenting lactose, see Figure 4. These cells reach stationary phase once the sugar sources have been depleted.

**Figure 3:**
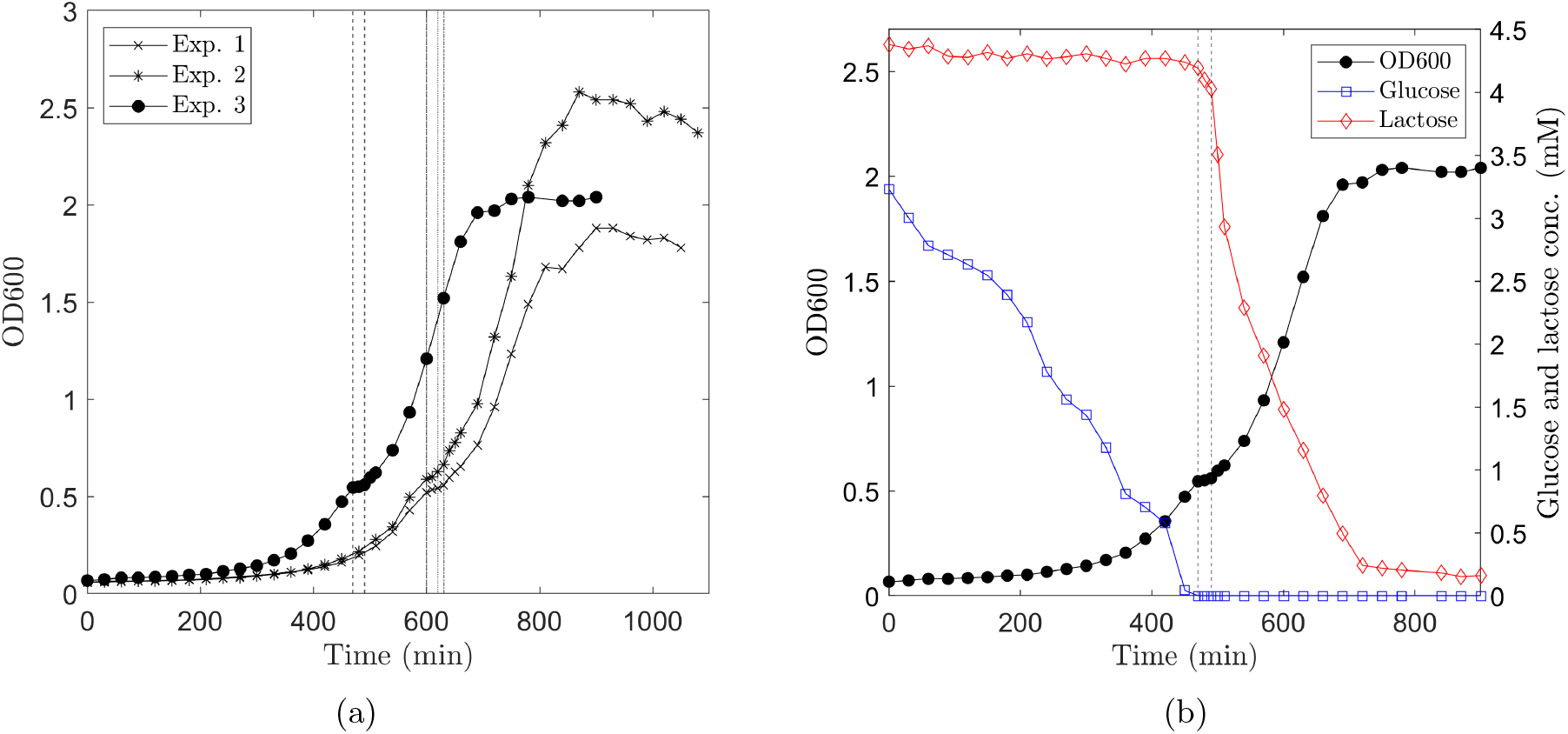
Mixed *E. coli* strains diauxic growth profile on glucose and lactose. (a) Growth curves of three independent biological replicates illustrating the transitions between initial lag phase, log phase, lag phase due to diauxic shift, log phase and then stationary phase as all sugars are depleted. (b) Glucose and Lactose concentrations relating to different parts of the growth curve for replicate experiment 3; black line and circles in (a), shown again here for completeness. Glucose (blue line, squares) is initially depleted by both strains (MV1717, MV1300) before a lag phase induced by the diauxic shift to lactose fermentation. Vertical dashed lines indicate the passage of diauxic shift.

**Figure 4:**
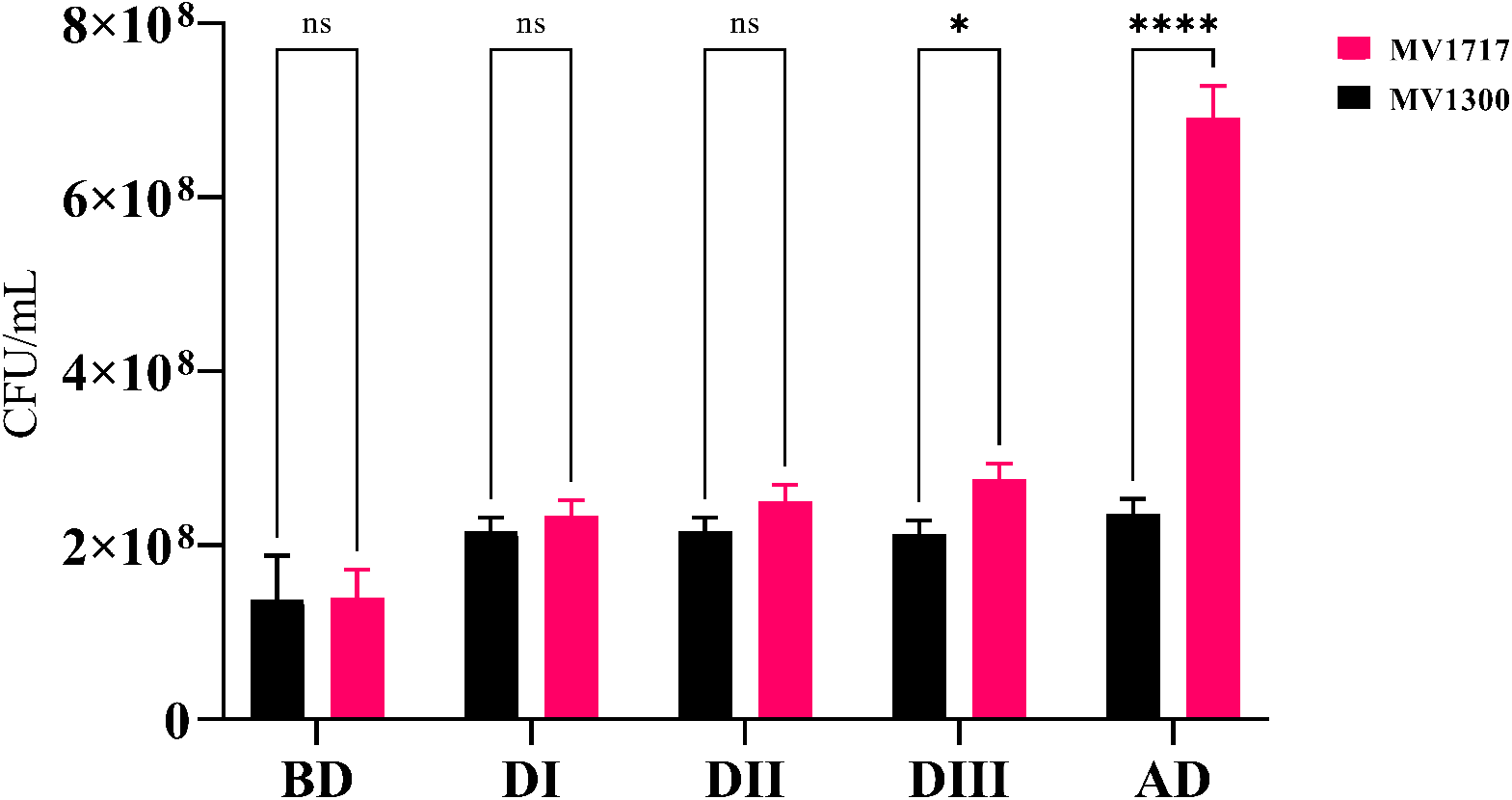
The enumeration of lactose (MV1717) and non-lactose (MV13000) fermenting strains on MacConkey agar. BD = samples in exponential growth before the diauxic shift (420 min); DI - DIII = samples during the diauxic shift (470, 480 and 490 min, respectively); AD = samples after the diauxic shift and during exponential growth (600 min). Bars represent the standard deviation of three replicates (n=3). The significance of differences was analysed by two-way ANOVA test (**** *P <* 0.0001; ns, not significant) and performed using GraphPad Prism software version 9.0.2 [27].

These results clearly demonstrate the lag phases associated with switching from rich to minimal media and glucose/lactose diauxie. Exponential growth is interrupted and then resumed as the cells switch metabolic pathways.

### 3.2 Modelling results

Full details of the governing equations and parameters used in the numerical simulations are given in Appendix F. The equations were solved in Matlab [28] using the solver ode15s.

#### 3.2.1 Comparison with experiment

We have two strains of *E. coli*, with concentrations *X*_1_ and *X*_2_, one of which, *X*_2_, cannot grow on lactose. Both strains are initially grown on a rich media (LB broth). The strains are then mixed in a 1 : 1 ratio and transferred to a minimal media containing a mixture of glucose (0.5g/L) and lactose (1.5g/L) as the only carbon source. Measurements of the concentrations of glucose, *S*_gl_, lactose, *S*_la_, and total biomass, *X*_1_ + *X*_2_, are taken at intervals from the point at which the strains are transferred to the minimal media, *t* = 0.

Results showing the predicted concentration of sugars and total biomass over time are shown as the solid lines in Figure 5(a) with the experimental data (shown as crosses) plotted for comparison. The mass fractions of the growth dependent parts of the proteome sectors are plotted in Figure 5(b) for both strains. The model predicts very slow initial biomass growth even though substrate is being taken up, which is in good agreement with the experimental data. The initial slow growth is due to the protein mass fractions being at non-optimum levels for growth on glucose in a minimal media. The strains, having previously been growing in a rich media, have a low level of anabolic A-sector proteins, Φ_*A*_ (see Figure 5(b)), resulting in slow growth. As the growth rate increases, the mass fractions reach optimum levels for glucose consumption (at around 6 hours). Just prior to exhaustion of glucose (at around 8 hours) the C-sector proteins are up-regulated and the R-sector and A-sector proteins are down-regulated in response to carbon deficiency. On depletion of glucose, strain 2 stops growing. The protein mass fractions of strain 1 are at non-optimal levels for lactose consumption and its growth slows, the lag phase. We measure the lag duration as the period when growth rate has dropped below 50% of the maximum on glucose. The model predicts the lag-phase to occur between 462 and 483 minutes in good agreement with the experimentally observed lag phase between 470 and 490 minutes. (The accuracy of determining the lag-phase duration from the data is obviously constrained by the frequency of measurements, in this case every 10 minutes).

**Figure 5:**
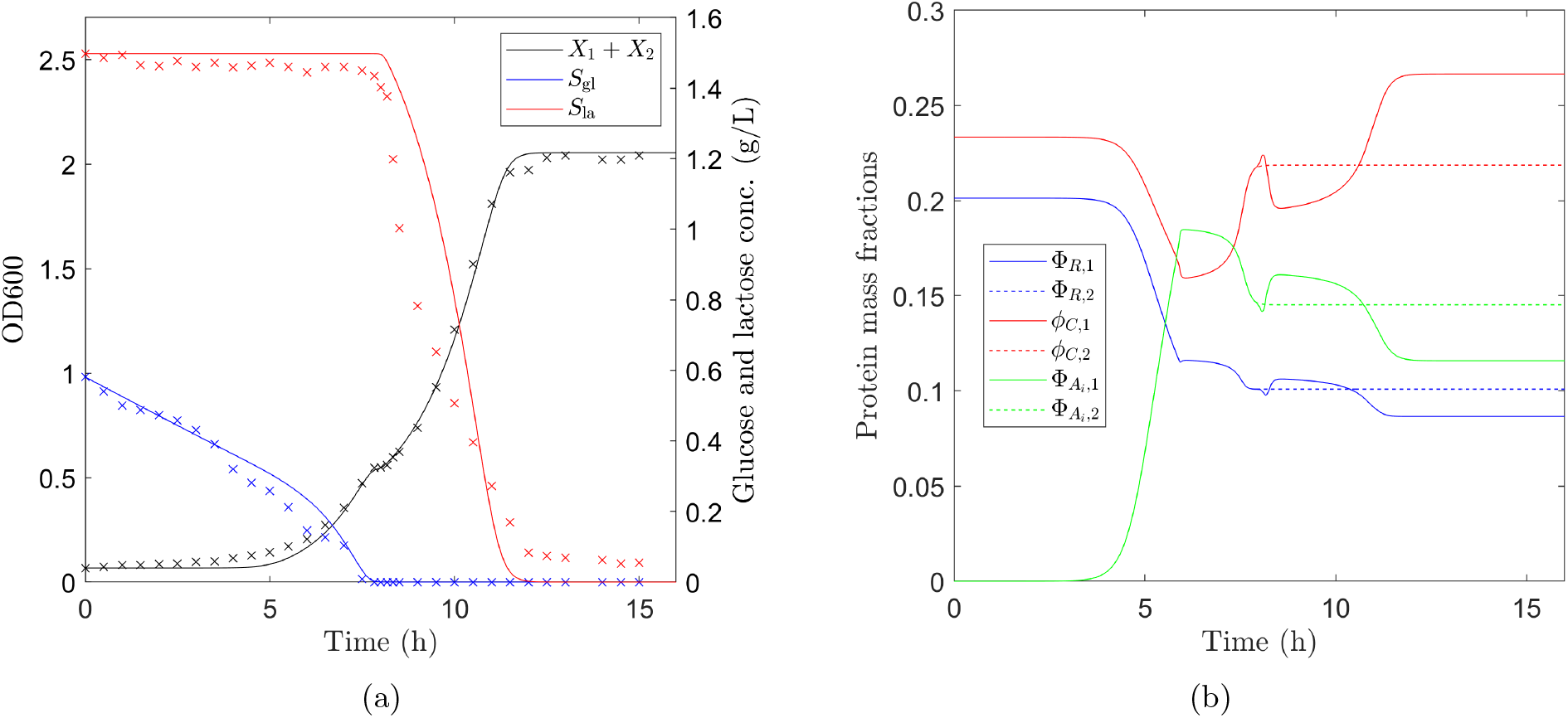
Mixed *E. coli* strains diauxic growth profile on glucose and lactose. (a) Solid lines show numerical predictions of glucose (blue) and lactose (red) concentrations and growth curve (black). Experimental data (Experiment 3 in Figure 3) are shown as crosses. The growth curve is the sum of biomass of both strains, *X*_1_ + *X*_2_. The model predicts a sequence of regimes. Initially there is very slow biomass growth even though substrate is being taken up, which is in good agreement with the experimental data. The diauxic shift can clearly be seen in the predicted growth curve at around 8 hours, in agreement with experiment. (b) Mass fractions of R-sector proteins (blue), C-sector proteins (red) and A-sector proteins (green) for strain 1 (solid lines) and strain 2 (dashed lines). The initial low level of A-sector proteins results in slow growth and, as protein production is proportional to growth rate, slow change in protein mass fractions. As the growth rate increases the mass fractions quickly reach optimum levels for glucose consumption (at around 6 hours). Just prior to exhaustion of glucose (at around 8 hours) the C-sector proteins are up-regulated and the R-sector and A-sector proteins are down-regulated in response to carbon deficiency. Strain 2 stops growing when glucose is depleted and its protein mass fractions stop changing. Strain 1 begins to consume lactose and readjusts its protein mass fractions to levels optimum for lactose consumption. As lactose levels decrease the C-sector proteins are again up-regulated and the R-sector and A-sector proteins down-regulated in response to carbon deficiency.

As strain 1 begins to consume lactose its protein mass fractions adjust to optimise lactose consumption. As lactose levels decrease the C-sector proteins are again up-regulated and the R-sector and A-sector proteins downregulated in response to carbon deficiency. This response to decreasing substrate levels is consistent with the response to a drop in nutrient quality [14]. However, as a microorganism enters the stationary phase different proteins, required for survival in nutrient deprived conditions, must be expressed [29]. We do not consider this in our model as we are primarily focused on describing the lag and log phases. This may explain the differences between model predictions of lactose concentration (zero at 12 hours) and experimental data (lactose not fully depleted after 12 hours).

It can be seen from Figure 5(a) that our description captures all principal features of the non-trivial growth curve of *E. coli* glucose-lactose diauxie. The lag-phase and diauxic shift are reproduced accurately using our rather fundamental model with minimal fitting and without the need for introducing an artificial lag parameter. All phases of growth, lag, log and diauxic shift, are determined from the structure of the microorganism’s proteome.

We now present the results of a sensitivity analysis looking at how changes to the fitted parameters affect the model predictions.

#### 3.2.2 Sensitivity analysis

The parameters *Y*_gl_, *Y*_la_, Φ _𝒢,0_ and *K*_*L*_ were determined by least-squares curve fitting to give a best fit to our experimental data. The best fit values are shown in Table 5 of Appendix F. Fitting was performed using the Matlab function lsqcurvefit [28] (for which the summed square of residuals for the fitted growth curve was equal to 0.0376).

Figure 6 shows numerical solutions for the total biomass for different values of the fitted parameters (experimental data for comparison shown as crosses). In all Figures the predicted growth curve using the best fit parameters is shown in red. In each subfigure one parameter is varied while all other parameters are fixed (to the values given in Tables 4 and 5 in Appendix F). As expected, changing the yields on glucose and lactose alters the final combined biomass concentration of the two strains; a lower yield value giving lower final biomass concentration and vice versa (see Figures 6(a) and 6(b)). Changing the minimum mass fraction of the key anabolic protein, Φ_*𝒢*,0_, alters the length of the initial lag phase and the final combined biomass concentration (Figure 6(c)). The lower the value of Φ_*𝒢*,0_ the slower the initial growth rate, increasing the time taken for the protein mass fractions to reach their optimum levels thus increasing the length of initial lag phase. The final biomass yield is also less for smaller Φ_*𝒢*,0_. The constant *K*_*L*_ determines when lactose starts to be consumed by strain 1 via the function

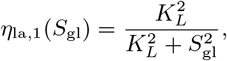

(note that strain 2 never consumes lactose). For values of *S*_gl_ ≫ *K*_*L*_ we have *η*_la,1_ ≪ 1 and lactose is not consumed. Only when glucose levels drop so that *S*_gl_ ≈ *K*_*L*_ will strain 1 begin to consume lactose. Reducing *K*_*L*_ increases the length of the diauxic lag phase as glucose levels must reach a lower value before lactose begins to be consumed (Figure 6(d)). If on the other hand *K*_*L*_ is increased the lag phase will shorten, with the extreme case *K*_*L*_ → ∞ (*η*_la,1_ = 1) removing the diauxic lag phase completely (glucose and lactose are consumed simultaneously). The final total biomass concentration as shown in Figure 6(d) is very slightly less for larger values of *K*_*L*_ suggesting that a diauxic shift may be beneficial for growth.

**Figure 6:**
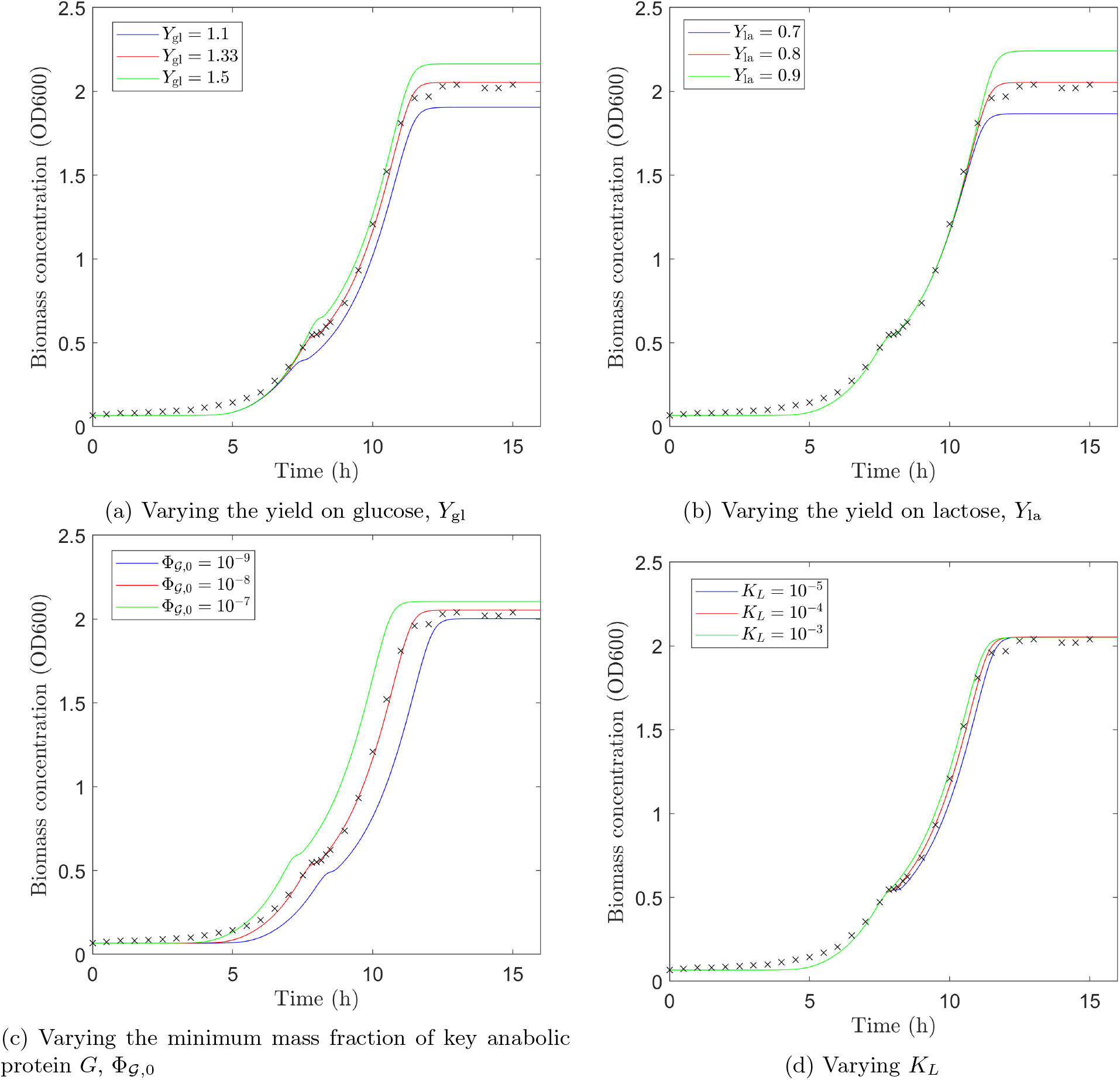
Comparison of numerical solutions when values of fitted parameters are varied (experimental data shown as black crosses). (a) The fitted value, *Y*_gl_ = 1.33, is shown in red. Reducing the yield (*Y*_gl_ = 1.1 shown in blue) decreases the biomass concentration. Conversely, increasing the yield (*Y*_gl_ = 1.5 shown in green) increases the biomass concentration. (b) The fitted value, *Y*_la_ = 0.8, is shown in red. Reducing the yield (*Y*_la_ = 0.7 shown in blue) decreases the biomass concentration post diauxic shift. Conversely, increasing the yield (*Y*_la_ = 0.9 shown in green) increases the biomass concentration post diauxic shift. (c) The fitted value, Φ_𝒢,0_ = 10^−8^, is shown in red. Reducing Φ_𝒢,0_ (Φ_𝒢,0_ = 10^−9^ shown in blue) reduces the initial growth rate, increasing the time taken for the protein mass fractions to reach their optimum levels thus increasing the length of initial lag phase. The final biomass yield is also reduced. Conversely, increasing Φ_𝒢,0_ (Φ _𝒢,0_ = 10^−7^ shown in green) increases the initial growth rate, shortens the initial lag phase and increases final biomass concentration. (d) The fitted value, *K*_*L*_ = 10^−4^, is shown in red. Reducing *K*_*L*_ (*K*_*L*_ = 10^−5^ shown in blue) means that the glucose level must reach a lower value before lactose can be consumed. This increases the length of the diauxic lag phase, though for the case shown this is a very small increase. Conversely, increasing *K*_*L*_ (*K*_*L*_ = 10^−3^ shown in green) means that lactose can begin to be consumed at higher glucose levels which shortens the diauxic lag phase (again this is a small effect for the case shown). The qualitative behaviour of the growth curve is similar for all cases shown.

#### 3.2.3 Applying the model to investigate different growth strategies

When resources are scarce, a strain that can outgrow its competitors will have an advantage [12]. In the following we compare two strains with the same initial biomass, hence we define the ‘better’ growth strategy as belonging to the strain with a higher final biomass concentration.

To examine whether diauxie is beneficial for growth we introduce a theoretical strain, *X*_3_, which does not exhibit diauxie, consuming glucose and lactose simultaneously, so that *η*_la,3_ = 1 (all other growth parameters are assumed to be the same as for *X*_1_ given in Tables 4 and 5). We use our parameterised model to predict growth curves when only *X*_1_ is present and then when only *X*_3_ is present. Initial conditions for both strains are identical and equal to those used in the simulation described in Section 3.2.1 (values given in Appendix F). The results, shown in Figure 7(a), show that the final biomass concentration is higher for *X*_1_ than *X*_3_: in this case, when only one strain is present, it is beneficial to grow diauxically. Strain 1 blocks the uptake of lactose when glucose is present, using all of the cells’ resources to metabolise glucose. Strain 3 cells must share their resources to break down glucose and lactose simultaneously, reducing the efficiency of glucose uptake. If however, we look at the case where strain 1 and strain 3 are competing for resources (essentially replacing *X*_2_ with *X*_3_ in our original model) we find that *X*_3_ outgrows *X*_1_, growth curves are shown in Figure 7(b). The advantage strain 3 has on depletion of glucose, by having no pause in growth, no diauxic shift, outweighs the initial inefficient glucose uptake.

**Figure 7:**
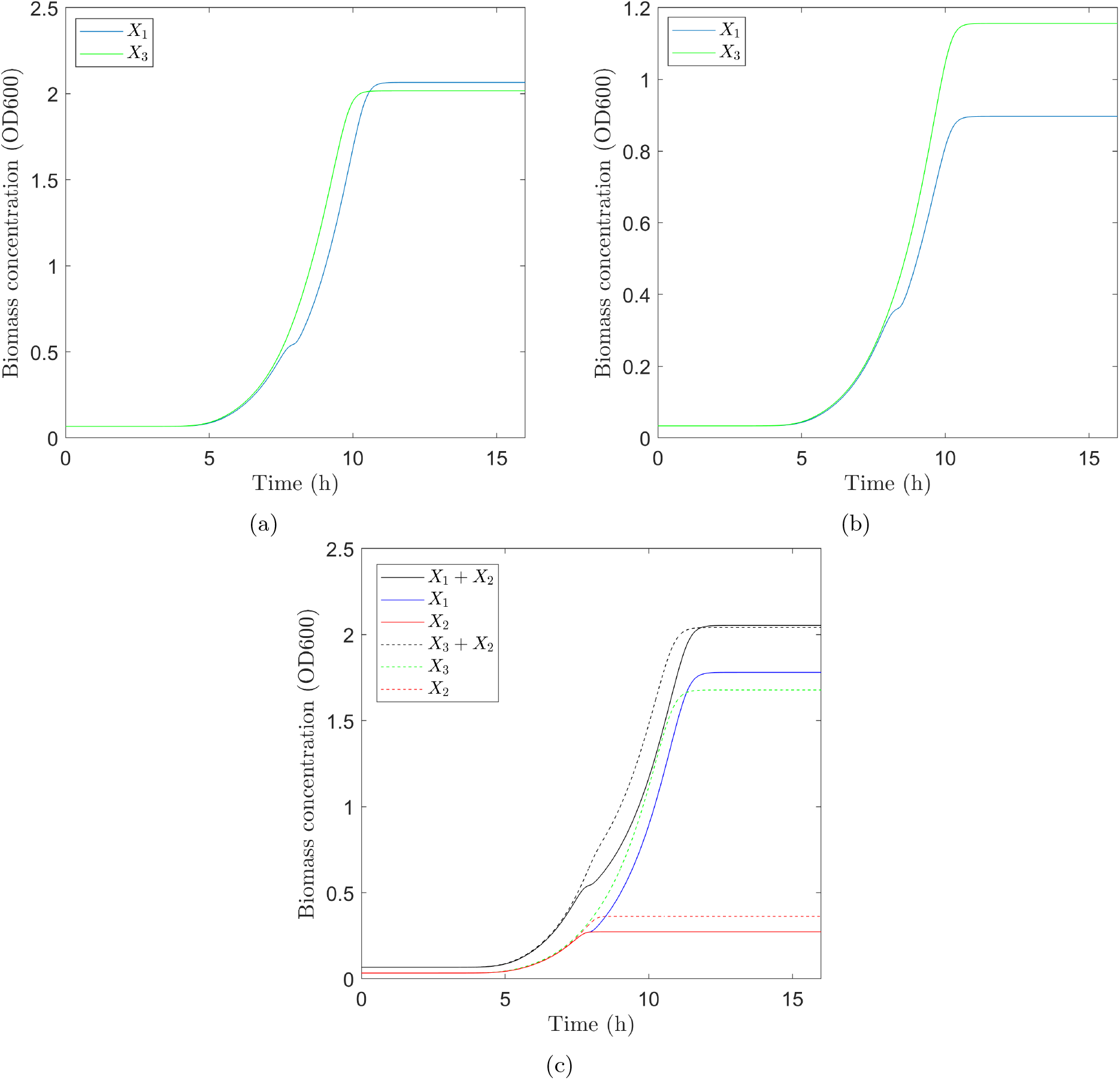
Growth curves for *E. coli* strains growing on a glucose/lactose mixture. (a) Growth curves from two simulations with only one strain present (no competition): diauxic strain *X*_1_ (blue); and non-diauxic strain *X*_3_ (green). The strain exhibiting diauxie, *X*_1_ has a slightly higher final biomass as it is able to consume glucose efficiently in the presence of lactose. (b) Growth curves from one simulation where two strains *X*_1_ (blue) and *X*_3_ (green) are competing for resources. The non-diauxic strain, *X*_3_, out-competes the diauxic strain, *X*_1_: in this particular case the advantage *X*_3_ has over *X*_1_ in being able to consume lactose efficiently immediately on exhaustion of glucose outweighs the disadvantage of its initial inefficient consumption of glucose. (c) Growth curves from two simulations: *X*_1_ and *X*_2_ competing for resources (solid lines); and *X*_3_ and *X*_2_ competing for resources (dashed lines). The mutant strain which cannot consume lactose, *X*_2_, performs better in competition with the non-diauxic strain *X*_3_ due to the inefficient consumption of glucose by *X*_3_.

We also re-ran the mixed strain parameterised model with *X*_3_ replacing *X*_1_ and compared the growth curve to our original fitted solution. Figure 7(c) shows the growth curves for the strains, combined (*X*_1_ + *X*_2_ and *X*_3_ + *X*_2_) and separately in these two cases. Although the final combined biomass concentrations are similar in both cases we see that *X*_1_ performs comparatively better than *X*_3_, with *X*_2_ performing better when growing with *X*_3_ than when *X*_1_ is present. Strain 1 can consume glucose as efficiently as strain 2, with the strains each consuming half of the total glucose. However, strain 2 out-competes strain 3, which is inefficiently simultaneously consuming glucose and lactose, consuming more than half of the total glucose and therefore growing more than when competing against strain 1.

We find that when a single strain of *E. coli* is growing on a mixture of glucose and lactose, it is better to consume the two sugars sequentially, however, when strains are competing for resources it is not necessarily beneficial for a strain to grow diauxically.

#### 3.2.4 Diauxic growth vs simultaneous consumption in a competitive environment

Diauxic growth is usually regarded as a process by which a microorganism maximises growth, however, during the diauxic lag phase there is a significant loss of growth. There is a trade-off between consuming the preferred sugar efficiently, maximising the microorganism’s long-term growth, and lost growth during the switch as the microorganism adjusts to using the secondary sugar. In a competitive environment are there conditions under which diauxic behaviour is an advantage and others which favour simultaneous consumption of resources?

Studies have shown that strains of bacteria [30] and yeast [31] can evolve to have differing lengths of diauxic shift. When a microorganism is subject to frequent changes in environment the diauxic lag will evolve to be short, whereas in a stable environment the lag phase will be longer [3].

Using our parametrised model we investigate the behaviour of two strains of *E. coli, X*_1_, exhibiting a diauxic shift, with *K*_*L*_ = 10^−4^, and *X*_3_, with no diauxic shift, *K*_*L*_ → ∞. This is essentially the same as the simulation shown in Figure 7(b), however we have changed the initial conditions to remove the initial lag phase, so that we are focusing on growth on glucose/lactose. We assume that the strains are mixed in a 1 : 1 ratio, *X*_1_ = *X*_3_ = 0.001, and that the initial protein mass fractions are the same, 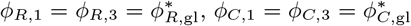. Results showing the ratio of biomass concentrations of the two strains, *X*_1_*/X*_3_, are shown in Figure 8.

**Figure 8:**
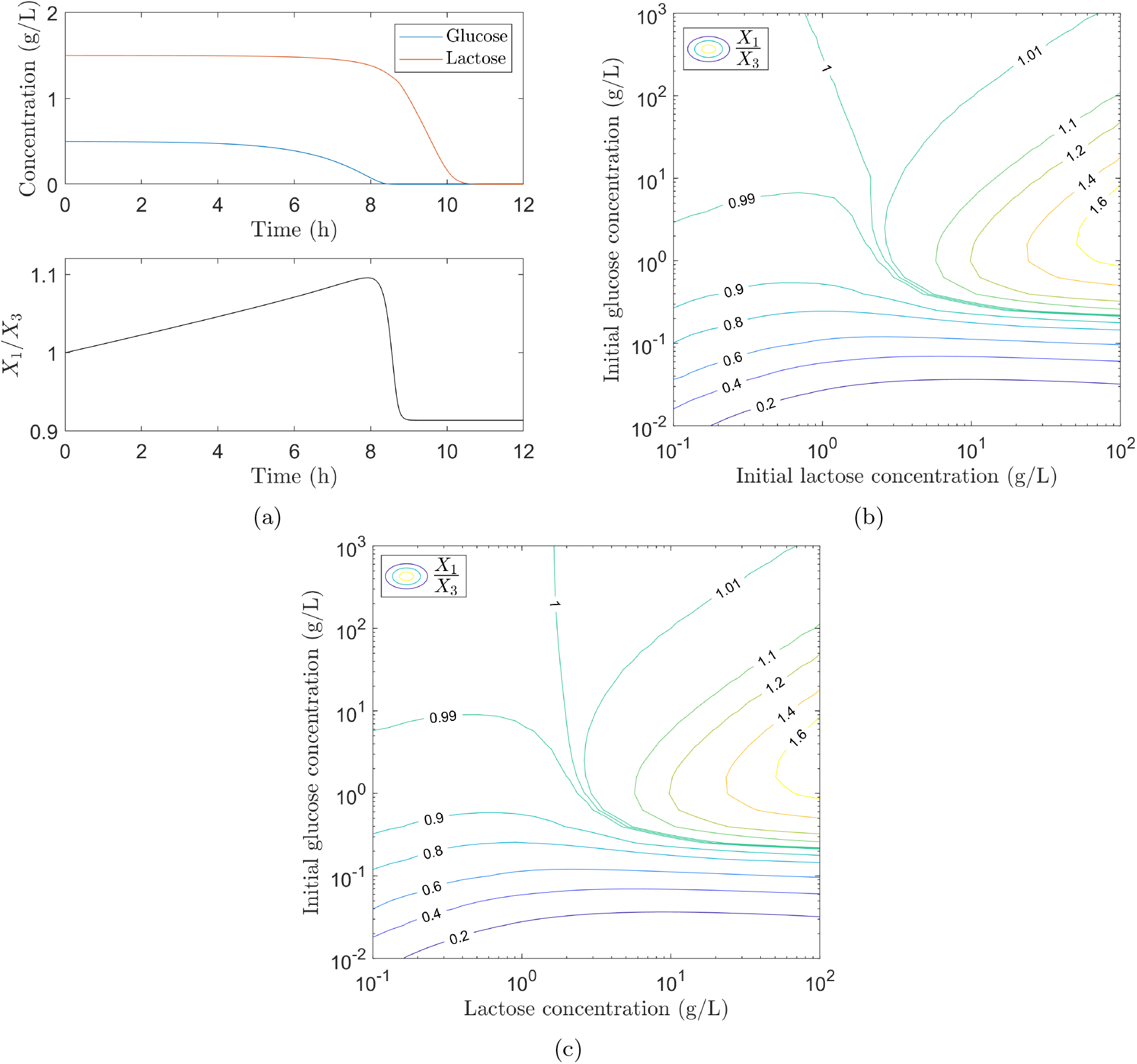
Comparison of two strains of *E. coli, X*_1_ which exhibits diauxie and *X*_3_ which has no diauxic shift, growing in a glucose/lactose mixture. (a) Upper figure shows concentration profiles for glucose (blue) and lactose (red) when initial concentrations are *S*_gl_ = 0.5g/L and *S*_la_ = 1.5g/L. The lower figure shows the ratio of the corresponding biomass concentrations *X*_1_/*X*_3_ (black). Initially *X*_1_ outperforms *X*_3_ (*X*_1_*/X*_3_ > 1) as it efficiently consumes only the preferred sugar glucose. As *X*_1_ enters the diauxic lag phase *X*_3_ continues to grow, as it is always able to consume lactose, quickly taking over from *X*_1_ and overall *X*_3_ outperforms *X*_1_ (the final ratio *X*_1_*/X*_3_ < 1). (b) Results showing final biomass ratio *X*_1_*/X*_3_ from simulations with varying different initial concentrations of glucose and lactose. For small initial concentrations of glucose and lactose *X*_3_ performs better (*X*_1_*/X*_3_ < 1): the advantage of being able to consume lactose efficiently immediately after the glucose is depleted outweighs the disadvantage of consuming lactose whilst the preferred sugar glucose is still available. Conversely, for larger initial glucose and lactose concentrations *X*_1_ performs better (*X*_1_*/X*_3_ > 1): the advantage of initial efficient consumption of the preferred sugar glucose outweighs the disadvantage of being unable to consume lactose efficiently immediately after the glucose is depleted. (c) Similar results to those shown in (b) are obtained when the lactose concentration is held fixed and only the glucose is depleted: *X*_3_ performs better at low sugar concentrations and *X*_1_ performs better at higher sugar concentrations.

Figure 8(a) shows results for the case where initial concentrations of glucose and lactose are *S*_gl_ = 0.5g/L and *S*_la_ = 1.5g/L respectively. The diauxic strain, *X*_1_, initially performs better than *X*_3_ (*X*_1_*/X*_3_ > 1): *X*_1_ is consuming only glucose but is doing so efficiently; *X*_3_ is consuming both glucose and lactose but at a reduced efficiency, hence a reduced growth rate. As the glucose is depleted *X*_1_ enters the diauxic lag phase and its growth rate drops dramatically. *X*_3_ continues to grow, as it is always able to consume lactose, quickly taking over and overall *X*_3_ outperforms *X*_1_ (the final ratio *X*_1_*/X*_3_ < 1).

This simulation was repeated for a range of different initial concentrations of glucose and lactose and for each simulation the final ratio *X*_1_*/X*_3_ was calculated. From the results, shown in Figure 8(b), we see that for small initial values of *S*_gl_ and *S*_la_ the non-diauxic strain will perform better *X*_1_*/X*_3_ < 1: the advantage of being able to consume lactose efficiently immediately after the glucose is depleted outweighs the disadvantage of consuming lactose whilst the preferred sugar glucose is still available. Conversely, for larger initial values of *S*_gl_ and *S*_la_ the diauxic strain performs better *X*_1_*/X*_3_ > 1: the advantage of initial efficient consumption of the preferred sugar glucose outweighs the disadvantage of being unable to consume lactose efficiently immediately after the glucose is depleted. Similar results are obtained when the lactose concentration is held fixed and only the glucose is depleted, see Figure 8(c).

When the concentrations of both sugars are kept constant the ratio *X*_1_*/X*_3_ will not tend to a fixed value as *t* → ∞ so we cannot use this to compare the strains in different conditions as above. However, when *S*_gl_ and *S*_la_ are fixed the growth rate, given by equation (2.15), is constant: *f*_gl_ and *f*_la_ only depend on substrate concentration and are therefore constant, and the mass fractions in constant conditions take their optimum values. The optimum values of the mass fractions satisfy equations (D.2), which are derived in Appendix D. Solving equations (D.2) for the mass fractions and substituting into equation (2.15) gives the growth rate as

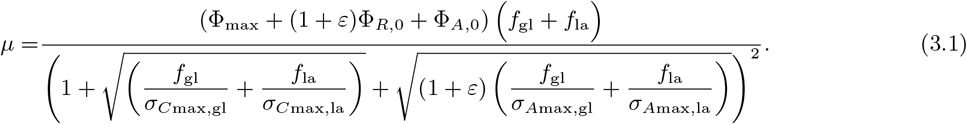

This equation was used to calculate the growth rate for the diauxic strain, *μ*_1_, and non-diauxic strain, *μ*_3_, for a range of values of *S*_gl_ and *S*_la_. The values obtained are shown in Figures 9(a) and 9(b) for *μ*_1_ and *μ*_3_, respectively. For both strains the growth rate never exceeds the value for maximum growth rate on glucose, 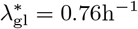 in our model. For *S*_la_ ≫ *S*_gl_ the growth rate approaches the value set for maximum growth rate on lactose, 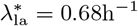 in our model.

**Figure 9:**
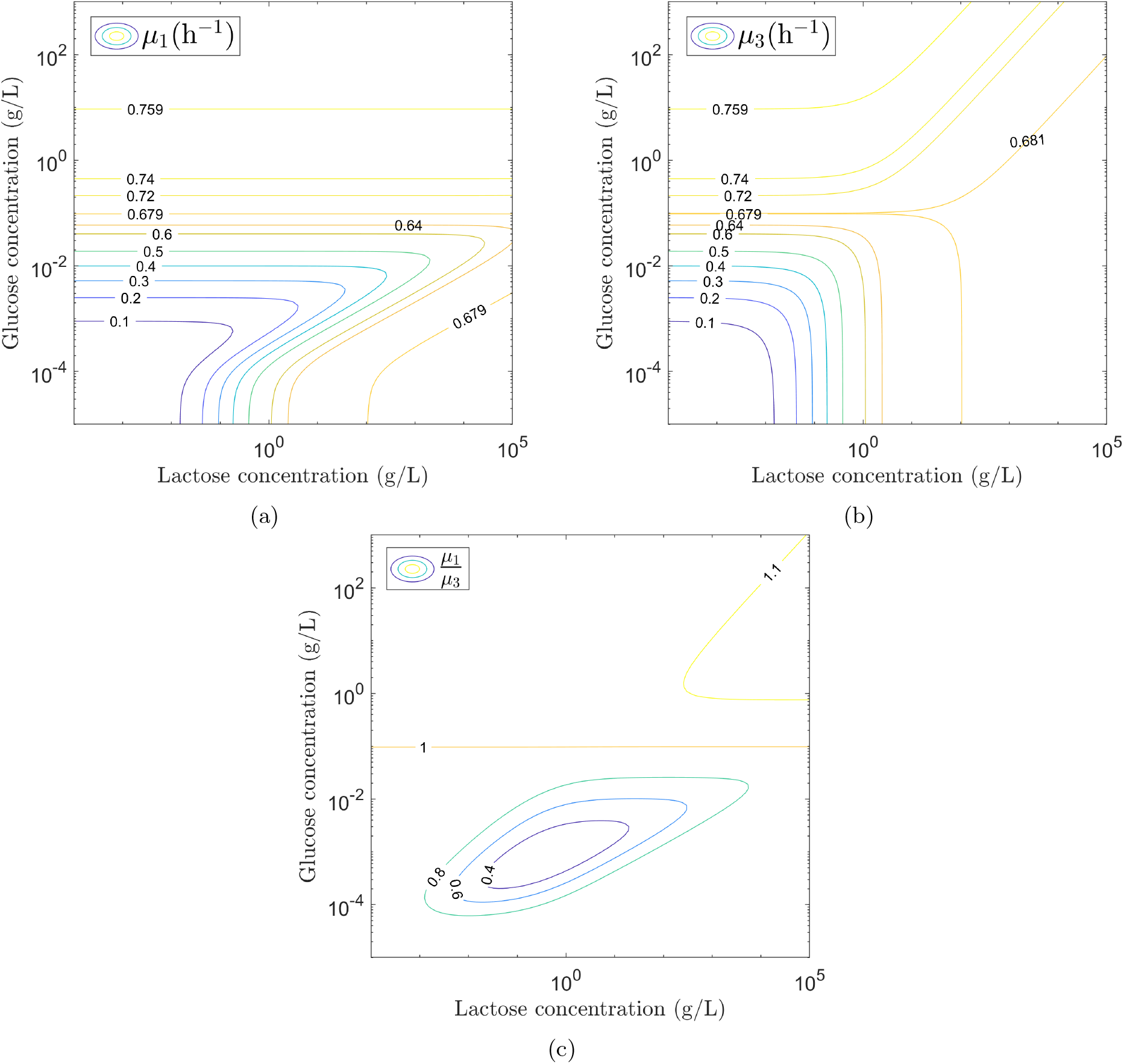
Growth rates of diauxic and non-diauxic *E. coli* strains growing in a glucose/lactose mixture under constant conditions. For each value of *S*_gl_ and *S*_la_ the growth rate of each strain is calculated using equation (3.1). Note that in order gain a fuller picture of the shape of the growth rate profiles the simulations were carried out for unrealistic *S*_la_ values (greater than the solubility of lactose). (a) Growth rate, *μ*_1_, of diauxic strain. *μ*_1_ never exceeds the value for maximum growth rate on glucose, 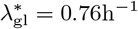, and for *S*_gl_ ≥ *O*(1) we have 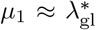. For *S*_la_ ≫ *S*_gl_ we have 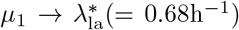. (b) Growth rate, *μ*_3_, of non-diauxic strain. As for the diauxic strain, *μ*_3_ never exceeds the value for maximum growth rate on glucose, 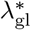, and for 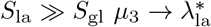. In this case, however, when 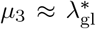 only at low lactose concentrations and as *S*_la_ increases *μ*_3_ decreases. (c) Contour plot of the ratio of the growth rates *μ*_1_*/μ*_3_. The growth rate of the diauxic strain is greater than that of the non-diauxic strain (*μ*_1_*/μ*_3_ > 1) if the glucose concentration is above a threshold value, *S*_gl,T_ ≈ 0.1. For *S*_gl_ > *S*_gl,T_ the diauxic strain will always outcompete the non-diauxic strain.

For constant growth rates, *μ*_1_ and *μ*_3_, equation (2.22d) for the biomass can be integrated to give

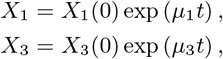

where *X*_1_(0) and *X*_3_(0) are the initial concentrations of strains 1 and 3 respectively. The ratio of strain concentrations is therefore *X*_1_*/X*_3_ = exp ((*μ*_1_ − *μ*_3_))*t*), as we have assumed that *X*_1_(0) = *X*_3_(0). This is greater than 1 for *μ*_1_ > *μ*_3_ and less than 1 for *μ*_1_ *< μ*_3_ so if the ratio *μ*_1_*/μ*_3_ > 1 it follows that *X*_1_ > *X*_3_ and the diauxic strain outperforms the non-diauxic strain and vice versa. Figure 9(c) shows a contour plot of the ratio of the calculated growth rates, *μ*_1_/*μ*_3_. The growth rate, *μ*_1_, of the diauxic strain is greater than that of the non-diauxic strain, *μ*_3_, except when the glucose concentration is below a threshold value, *S*_gl,T_ (for our simulations we have *S*_gl,T_ ≈ 0.1). For *S*_gl_ *< S*_gl,T_ the growth rate on glucose drops as *S*_gl_ ≈ *K*_*S*,gl_: both strains are consuming glucose inefficiently. As *X*_3_ is able to consume lactose as well as glucose its growth rate, *μ*_3_ is higher than *μ*_1_. For *S*_gl_ > *S*_gl,T_, the diauxic strain outperforms the non-diauxic strain: strain *X*_1_ only consumes glucose and all of the cell’s resources are optimally used for metabolising glucose, conversely, strain *X*_3_ will always be simultaneously consuming glucose and lactose, its limited resources shared between breaking down the two sugars resulting in a reduced growth rate. Note that when there is no lactose at all in the system the performance of the two strains is the same.

For constant sugar concentrations we have a stable growth environment. Our results show that, assuming there is a reasonable amount of preferred sugar available (*S*_gl_ > *S*_gl,T_), in a stable environment a strain exhibiting diauxie will out-perform a non-diauxic strain. Further, from the results in Figures 8(b) and 8(c) the diauxic strain out-performs the non-diauxic strain at higher sugar concentrations. The growth environment changes on depletion of glucose/lactose so the higher the initial sugar concentrations the longer the time before conditions change. Therefore, the environment can be considered stable for a longer period as the initial concentrations of the two sugars, particularly glucose, increases. Thus, the results in Figures 8(b) and 8(c) show that as the environment becomes more stable a diauxic strain will out-perform a non-diauxic strain.

Our simulation results agree qualitatively with experimental evidence that bacteria [30] and yeast strains [31] with a long diauxic lag perform better in a stable environment and those with a short (or no) diauxic lag perform better in a changing environment.

## 4 Discussion

In this paper, we have formulated a coarse-grained mechanistic model describing the time evolution of biomass growth, substrate concentration and gene expression during carbon upshifts and downshifts. The model extends recent descriptions, incorporating proteome partitioning, flux-controlled regulation and optimal allocation of protein synthesis. Carbon influx is balanced with amino acid and protein synthesis fluxes via adjustments to the amino acid synthesis rate and average translation rate, the rates being determined by the size of pools of central precursors (including ketoacids and amino acids). Here, we recognise that the central precursors are limited by the innate capacity of a cell; the model includes a mechanistic functional response to limit the size of the precursor pools, ruling out physically unrealizable behaviour observed in earlier studies.

Phases of microorganism growth emerge from the dynamics, rather than being switched on/off at a particular time. The selective use of substrates, regulated by mechanisms such as CCR, is achieved by completely different methods in different microorganisms [6]. Accordingly, the exact mechanism underlying the inhibition of substrate uptake is not made explicit in our model, making it flexible and applicable to processes other than *E. coli* glucose-lactose diauxie, which we have focussed on. The switch to consuming the secondary substrate, controlled through functions *η*_*j*_, occurs when the concentration of the preferred substrate drops below a set value, *K*_*L*_. In this way we avoid having to artificially switch on the inferior carbon uptake system at a predetermined time as in other models [18, 32].

Furthermore, the regulation functions allocating protein synthesis are obtained by mathematically optimising the growth rate. Resource allocation in steady state conditions can be determined from fundamental growth laws relating protein levels to growth rate [11, 14, 18, 24]. In dynamic conditions, however, it remains unclear how protein synthesis is regulated. Erickson et al [18] construct regulation functions based on the steady state growth laws but these suffer from being undefined or negative during growth transitions. Therefore, we formulated an alternative description of protein allocation, that is valid during growth transitions, where the regulation functions are derived directly via mathematical optimization of the growth rate.

Employing our modelling approach, we found that phases of bacterial growth, including the lag phase and diauxic shift, emerged from the structure of the bacterial proteome. In particular, the deterministic model predicted the lag-phase and diauxic growth of *E. coli* on glucose and lactose. We found that the predicted diauxic shift occurs between 462 and 483 minutes, which is in good agreement with the experimentally observed times of 470 and 490 minutes. The primary focus of the current study was to describe lag and log-phase growth. Therefore, the transition to stationary phase is less well captured (as demonstrated by inconsistencies between predicted and measured lactose concentration post 12 hours). This could be addressed by taking account of the expression of proteins required for survival in nutrient deprived conditions [29].

Earlier dynamic resource allocation models have focused on predictions of growth rate/biomass and protein levels [17, 18]. However, these models are unable to capture the non-simple relationship between substrate uptake rate and growth rate observed experimentally during the lag phase. Substrate concentrations have been predicted in the rather large and complex modelling approach of Salvy and Hatzimanikatis [5]. However, we found that our much simpler coarse-grained model was sufficient to describe the time evolution of substrate concentrations in addition to biomass and protein levels, accurately replicating the observed relationship between substrate uptake and biomass growth during lag phase.

When a microorganism switches between carbon sources there is a trade off between optimising growth on the preferred substrate and being able to switch quickly when the primary source is depleted [3, 19]. We have shown that the lag phase observed when *E coli*. switches from a rich to a minimal media can be explained by a low level of a key anabolic protein causing a bottleneck in the metabolic flux pathway. This agrees with the conclusions of Basan et al [19] that lag phases are caused by metabolic bottlenecks.

Our investigation into the merits of different bacterial growth strategies finds that in a stable environment a strain exhibiting diauxie will always outcompete a non-diauxic strain (except at low glucose concentration, *S*_gl_ ≲ *K*_*S*,gl_).

The non-diauxic strain always has the lactose metabolism switched on meaning that metabolic resources must be shared between the breakdown of glucose and lactose. When both sugars are present this strain therefore operates at a reduced efficiency with a lower growth rate, however, when glucose is depleted the strain can immediately grow on lactose as lactose metabolism is always switched on. This behaviour is in agreement with results of Chu and Barnes [3] that premature activation of the secondary metabolism shortens the lag but causes costs to the cell thus reducing the growth rate on the preferred substrate.

Recent work has made clear that microorganisms living in changing environments do not always favour perfect catabolite repression [31, 32, 33]. New et al [32] found that although stringent catabolite repression seems favourable in relatively stable environments, less stringent regulation can increase fitness in variable conditions. To explore competition in a changing environment we ran simulations starting with a limited amount of sugar. The growth environment changes on depletion of glucose/lactose so the higher the initial sugar concentrations the longer the time before conditions change. The environment is therefore stable for a longer period as the initial concentrations of the two sugars, particularly glucose, increases. Our results show that a diauxic strain prevails when initial sugar concentrations are high but a non-diauxic strain, that switches immediately on depletion of glucose, dominates at low initial sugar concentrations. Our results compare favourably with the results of New et al [32] and other experimental evidence that bacteria [30] and yeast strains [31] with a long diauxic lag dominate in a stable environment and those with a short (or no) diauxic lag dominate in a changing environment.

This study adds to the rich body of work showing how microorganisms react to changing environments [5, 18, 32, 34, 35]. The range of applications of our modelling approach is large: the description can be generalised to model multiple different microorganisms, investigate competition between different species or strains and explore other growth strategies. The model can be adapted to predict the growth of many bacteria and yeasts that exhibit diauxie. More generally, the model provides a means to investigate and describe lag phase, the mechanisms for which, despite many years of research, are only just being revealed.

## Statements and Declarations

The authors have no competing interests to declare that are relevant to the content of this article.

## Data availability

All data generated or analysed during this study are included in this published article.

## Author contributions

All authors contributed to the study conception and design. Mathematical model construction and analysis were performed by Fiona Bate. Experimental material preparation, data collection and analysis were performed by Yumechris Amekan. The original draft of the manuscript was written by Fiona Bate and Yumechris Amekan and all authors reviewed and edited the manuscript. All authors read and approved the final manuscript.

## Funding

This work was supported by a Daphne Jackson Trust Fellowship funded by the University of York and Biotechnology and Biological Sciences Research Council; the Indonesia Endowment Fund for Education; and a Royal Society Industry Fellowship (IF160022).

## Acknowledgments

We would like to thank Luna Yuan for informative discussions.

## Appendix

### A. The values of the mass fractions during the log phase of growth

It has been shown experimentally that during the log phase of growth of bacterial cells, the rate of cell proliferation (the growth rate) and the expression levels of key proteins are linearly correlated [11, 14, 18]. In these experiments *E. coli* cells are grown on a range of different nutrients. Once the cells reach the log phase of growth measurements of protein expression levels and the growth rate are taken. In the following a star denotes the value of a variable during the log phase of growth.

#### A.1 R-sector proteins

Experimentally the RNA/protein ratio, *r*, which is a well established proxy for the ribosomal mass fraction [18], is measured rather than Φ_*R*_ itself. For cells growing exponentially under nutrient (e.g. carbon or nitrogen) limitation, *r* is linearly correlated with the growth rate [14] and we have the following bacterial growth law

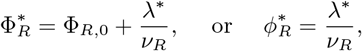

where *λ*^*^ is the growth rate of *E. coli* cells during log phase, Φ_*R*,0_ is the growth independent minimum level of Φ_*R*_ and *ν*_*R*_ is a constant [18]. Values of these parameters taken from the literature are shown in Table 3 (the relation Φ_*R*_ = 0.46 *r* was used to fit the data in [18]).

**Table 3:** Parameter values taken from the literature that are used in the equations defining the value of mass fractions during the log phase. These equations are described fully in the text.

| RNA/protein ratio (C-limitation)<br>$r^* = r_0 + \frac{\lambda^*}{\nu_R}$ <sup>c</sup> | | Ribosomal mass fraction (C-limitation)<br>$\Phi_R^* = \Phi_{R,0} + \frac{\lambda^*}{\nu_R}$ <sup>c</sup> | |
| --- | --- | --- | --- |
| $r_0$ | $\nu_R$ (h<sup>-1</sup>) | $\Phi_{R,0}$ | $\nu_R$ (h<sup>-1</sup>) |
| $0.076 \pm 0.005$ <sup>a</sup> | $5.3$ <sup>a</sup> | $0.049 \pm 0.02$ <sup>b</sup> | $11.02 \pm 0.44$ <sup>b</sup> |
| PLacZ expression (C-limitation)<br>$L^* = L_{\max} \left(1 - \frac{\lambda^*}{\lambda_L}\right)$ | | | |
| $L_{\max}$ (MU $\times 10^3$) | | $\lambda_L$ (h<sup>-1</sup>) | |
| $32 \pm 1$ <sup>a</sup> | | $1.2 \pm 0.0$ <sup>a</sup> | |
| $21.8 \pm 0.5$ <sup>b</sup> | | $1.17 \pm 0.05$ <sup>b</sup> | |

<sup>a</sup>Data from [11].
<sup>b</sup>Data from [18].
<sup>c</sup>The two expressions for the ribosomal bacterial growth law given in [11] and [18] respectively are interchangeable. In [18] it is presented in terms of the ribosomal mass fraction, although the RNA/protein ratio is measured, and the relation $\Phi_R = 0.46r$ is used to fit the model to data. Further details are given in sections A.1 and A.2.

#### A.2 C-sector proteins

The value of 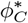 is determined by assuming proportionality to a reporter enzyme (P*LacZ* in [11] and [18]). Experimentally it has been shown [11, 14, 18] that when carbon is limiting growth we have

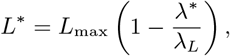

where *L*_max_, the maximum level of the reporter enzyme, *L*, and the constant *λ*_*L*_ are determined by fitting to experimental data. Values of these parameters taken from the literature are shown in Table 3.

The growth dependent part of the catabolic protein sector is assumed to be regulated as a whole [11, 24] so for each catabolic enzyme with mass fraction Φ_*E*_ we have

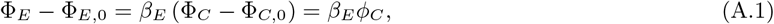

where Φ_*E*,0_ is the growth independent minimum level of the enzyme and *β*_*E*_ is a constant. For the reporter enzyme, *L*, we therefore have *L* − *L*_0_ = *β*_*L*_*ϕ*_*C*_, where *L*_0_ is the growth independent minimum level of reporter enzyme (the total protein level is independent of growth rate [11] so *L* is directly proportional to the mass fraction of reporter enzyme). When the catabolic sector is at its maximum this gives *L*_max_ − *L*_0_ = *β*_*L*_Φ_max_ and it follows that

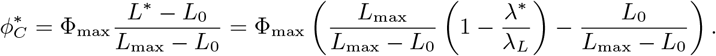

Experimental results show that *L*_max_ ≫ *L*_0_ [11, 18] so the above expression can be simplified by neglecting small terms to give

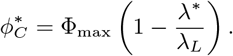

In deriving our model equations the mass fraction of a catabolic enzyme, Φ_*E*_, only appears as part of the fraction 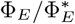 (see Appendix B). From equation A.1 we obtain

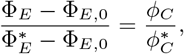

which can be simplified by assuming 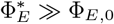 (as supported by experimental evidence [11]), to give

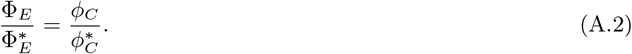

Using this we can express the governing equations of our model explicitly in terms of *ϕ*_*C*_.

#### A.3 A-sector proteins

The A-sector is assumed to be regulated as a whole [11, 24] so for an anabolic protein, 𝒢, with mass fraction Φ_*𝒢*_, we have

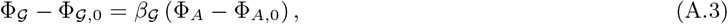

where Φ_*𝒢*,0_ is the growth independent minimum level of protein 𝒢 and

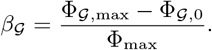

Rescaling the mass fraction so that 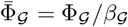, we can remove *β*_*𝒢*_ from equation (A.3) obtaining

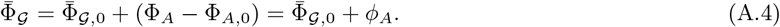

The value of *ϕ*_*A*_ can be determined from *ϕ*_*R*_ and *ϕ*_*C*_ : from equation (2.2) we have

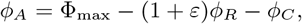

which yields

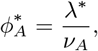

with

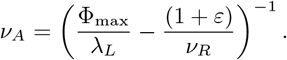

### B Substrate uptake - multiple substrate enzyme kinetics

Following Michaelis-Menten kinetics [36], multiple substrates with concentrations *S*_*j*_ are broken down by a key catabolic enzyme, *E*, in the following way

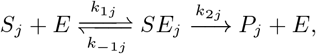

where *P*_*j*_ represents the product of the reaction. The reactions to form the enzyme-substrate complexes *SE*_*j*_ are assumed to be reversible with the rate of the forward reaction given by *k*_1*j*_ and the reverse reaction given by *k*_−1*j*_. The product forming reactions are not reversible with rate given by *k*_2*j*_. For the case when the presence of one substrate prevents the consumption of substrate 𝒥 (as occurs with glucose-lactose diauxie) the formation of the enzyme substrate complex will not occur for substrate 𝒥. To model this we write 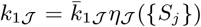 where 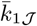 is the rate of the forward reaction when only substrate 𝒥 is present and *η*_𝒥_ ({*S*_*j*_}) depends on other substrates present in the system. Under conditions which favour the uptake of substrate *j* we have *η*_*j*_ = 1.

Applying the law of mass action we obtain the ordinary differential equations

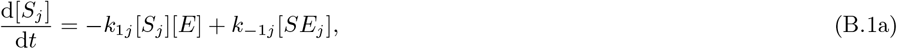

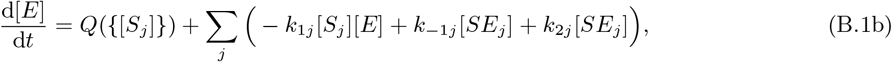

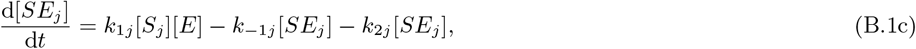

where *Q*({[*S*_*j*_]}) gives the rate of enzyme production and square brackets denote concentration. We assume that the enzyme-substrate complex is formed on a much faster timescale than product formation and that the concentration of complex does not change on the time-scale of product formation (quasi-steady-state assumption). We therefore set

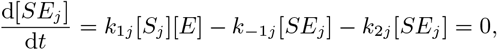

so that

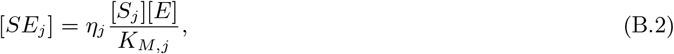

where

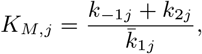

is the Michaelis constant for substrate *j*. We now introduce 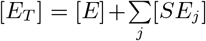 to be the total amount of catabolic enzyme present in the system (the free enzyme plus the enzyme in the substrate-enzyme complexes). Substituting in for [*SE*_*j*_] from equation (B.2) and rearranging gives

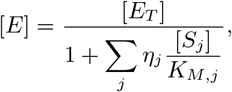

which on substitution back into equation (B.2) gives

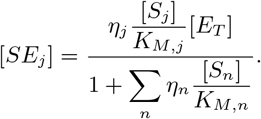

Equations (B.1) become

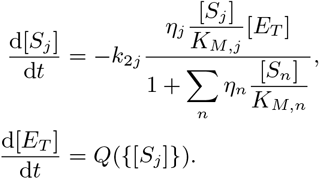

Writing [*E*_*T*_] = *p* Φ_*E*_ [*X*], where the constant *p* (introduced in Section 2.2.2) is the fraction of biomass that is protein, the substrate equations become

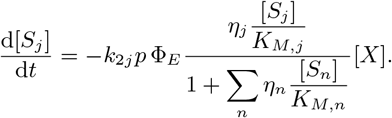

The maximum uptake rate, *k*_max,*j*_, occurs during the log-phase of growth when substrate *j* is the only substrate present and is in excess (*S*_*j*_ ≫ *K*_*M,j*_) so that

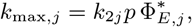

where 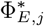 is the value of the mass fraction of enzyme during log-phase growth on substrate *j*. It follows that

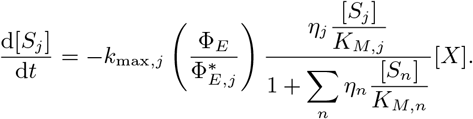

Using equation (A.2) we can write this in terms of the mass fraction of the total growth dependent catabolic sector,

*ϕ*_*C*_, giving

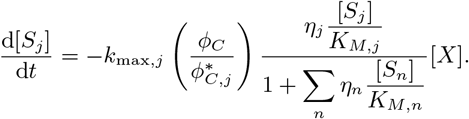

In the main text the brackets denoting concentration are dropped: substrate and biomass concentrations in the main text are denoted by *S*_*j*_ and *X* respectively.

### C Solving the flux balance equations to obtain the carbon influxes, *J*_*C,j*_

To keep the number of variables in the model to a minimum we can remove the explicit dependence of the carbon influxes, *J*_*C,j*_, on *P*. In this section we solve the flux balance equations to obtain *P*_*C,j*_ and *P*_*A,j*_, and hence *P*, only in terms of the substrate concentrations and protein mass fractions. We can then eliminate *P* from the equation for *J*_*C,j*_.

The flux balance equations, given in the main text in equation (2.11), are

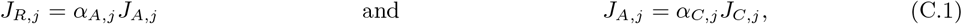

with the fluxes given by 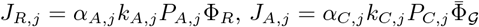 and, from equation (2.10),

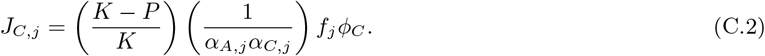

Substituting for *J*_*R,j*_ and *J*_*A,j*_ into the first of equations (C.1) and rearranging we obtain

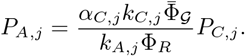

It follows, from equation (2.8), that the combined size of precursor and amino acid pools is given by

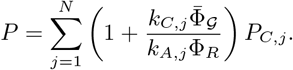

This expression for *P* can now be substituted into equation (C.2) from which we obtain

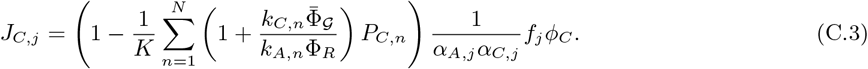

Substituting for *J*_*A,j*_ and *J*_*C,j*_ into the second of equations (C.1) and rearranging we obtain

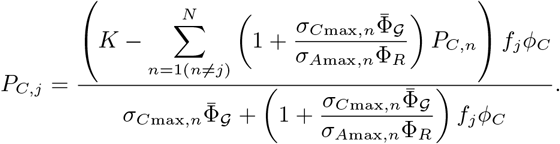

To simplify the notation we have introduced *σ*_*A*max,*n*_ = *α*_*A,n*_*α*_*C,n*_*k*_*A,n*_*K*, the maximum translation rate when only substrate *n* is being consumed. The definition of *σ*_*A*max,*n*_ follows from substituting the maximum possible value for *P*_*A,n*_ = *α*_*C,n*_*K* (which comes from equation (2.8) with *P* = *K, P*_*A,j*_ = 0 for *j* ≠ *n* and *P*_*C,j*_ = 0 ∀*j*) into our expression for the translation rate 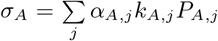. We similarly define *σ*_*C*max,*n*_ = *α*_*A,n*_*α*_*C,n*_*k*_*C,n*_*K* which is *α*_*A,n*_ times the maximum amino acid synthesis rate when only substrate *n* is being consumed. (Defining *σ*_*C*max,*n*_ as being *α*_*A,n*_ times the maximum amino acid synthesis rate instead of just the maximum amino acid synthesis rate further simplifies our notation.)

We now have each *P*_*C,j*_ in terms of all the other *P*_*C,n*_, a total of *N* equations for *N* unknowns. We solve the equations by first eliminating *P*_*C*,1_ from all the equations then *P*_*C*,2_, until we obtain an expression for *P*_*C,N*_ which does not reference any other *P*_*C,n*_. For *j* = 1 we have

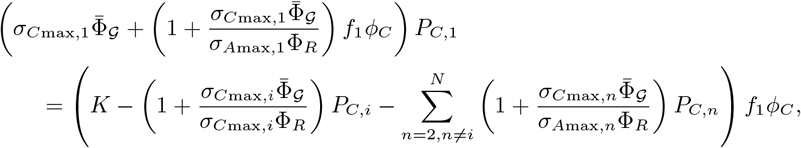

and for *j* = *i* ≥ 2 we have

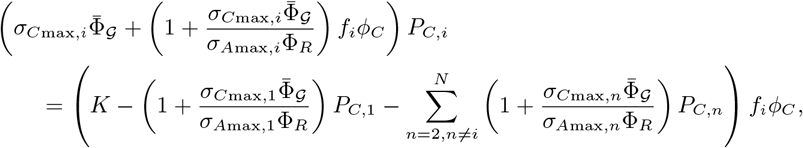

where we have written out explicitly terms in *P*_*C*,1_ and *P*_*C,i*_. Eliminating *P*_*C*,1_ from these equations we obtain

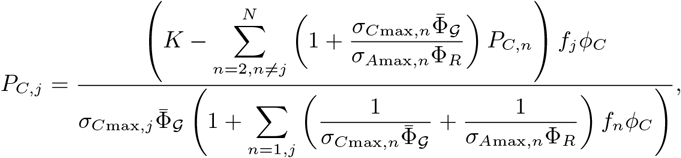

where 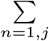 denotes the sum of the 1^st^ and *j*^th^ terms. Again, writing out explicitly terms in *j* = 2 and *j* = *i* ≥ 3 we have for *j* = 2

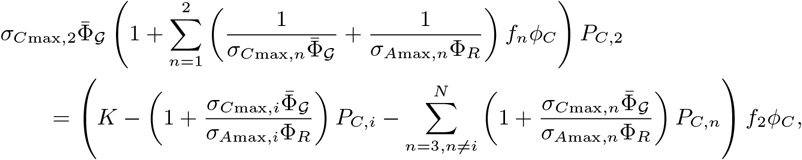

and for *j* = *i* ≥ 3

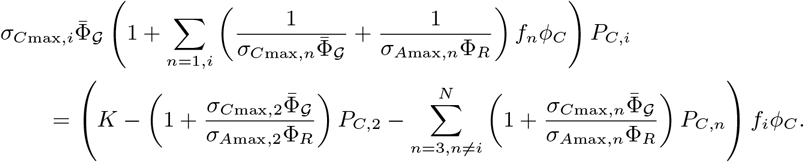

Eliminating *P*_*C*,2_ from these equations we obtain

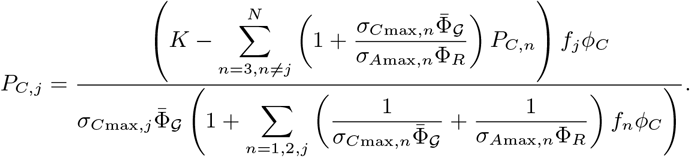

Carrying on in this way we obtain an expression for *P*_*C,N*_ as

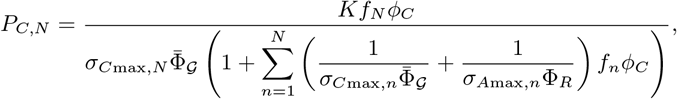

and in general we have

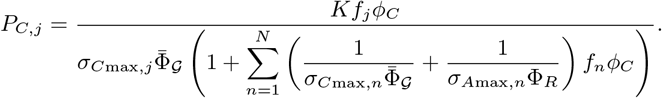

Substituting into equation (C.3) and rearranging we obtain

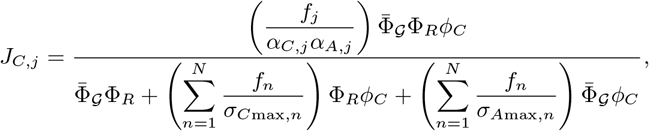

which is the carbon influx from substrate *j* in terms of only the substrate concentrations {*S*_*j*_} (through the substrate dependent functions {*f*_*j*_}) and the protein mass fractions Φ_*R*_, 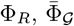 and *ϕC*.

### D Determining the unknown constants *σ*_*C*max,*j*_, *σ*_*A*max,*j*_ and *α*_*A,j*_*α*_*C,j*_*Y*_*C,j*,0_ in terms of experimentally measurable parameters

The expression for the growth rate, given in the main text by equation (2.15), is

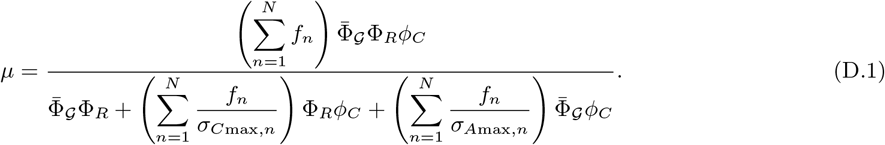

This contains the unknown constants *σ*_*C*max,*j*_, *σ*_*A*max,*j*_ and *α*_*A,j*_ *α*_*C,j*_ *Y*_*C,j*,0_, the latter combination of constants appearing in the definition of *f*_*j*_ (equation (2.9)). We cannot determine these constants directly from known experimental measurements so instead we relate them to measurable parameters such as growth rate, biomass yield and protein mass fraction. We do this by finding the maximum value of *μ* in terms of the protein mass fractions and assuming that during the log-phase of growth the growth rate is equal to this maximum value.

We optimise *μ* in terms of 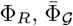 and *ϕ*_*C*_ using the method of Lagrangian multipliers with the constraint given by

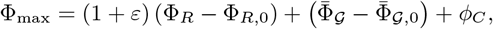

which is obtained from equation (2.2). Writing

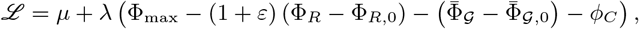

it follows that

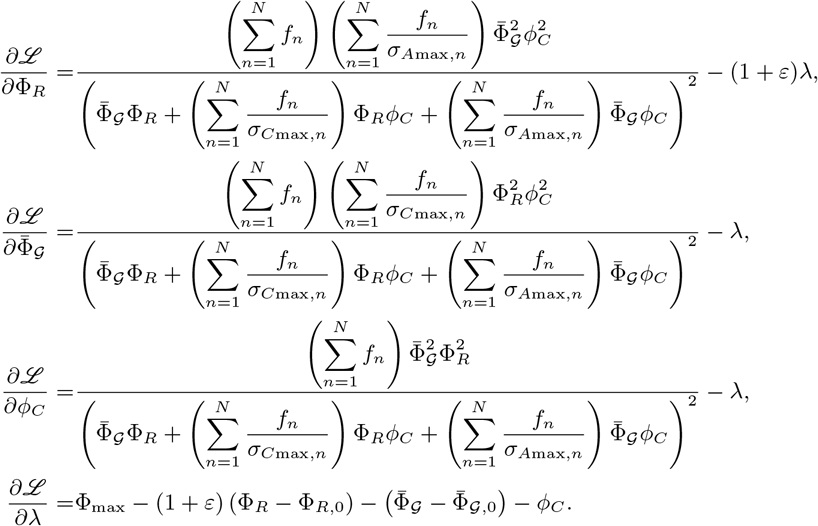

Setting all partial derivatives equal to zero and eliminating *λ* we obtain

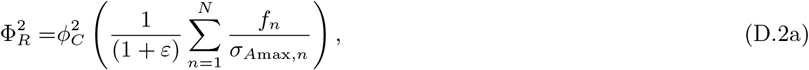

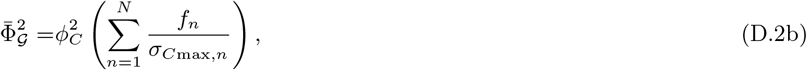

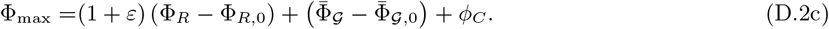

We denote experimental measurements of biomass yield and growth rate on a single substrate *S*_*j*_ during the logphase of growth by 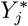 and 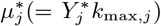 and mass fraction values by 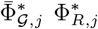 and 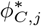. The values of the mass fractions during the log-phase are calculated from equations (2.3). Now, assuming that during log-phase the growth rate, 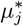, takes its maximum value, the calculated values for the mass fractions must satisfy equation (D.2) and it follows from equations (D.2a) and (D.2b) that

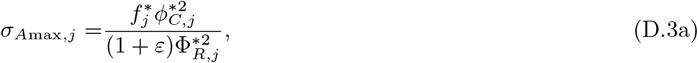

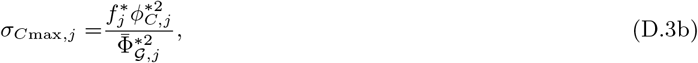

where 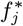 is the value of the function *f*_*j*_ during the log-phase of growth on substrate *j*. From equation (D.1) we have

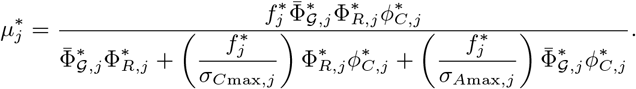

which after elimninating *σ*_*A*max,*j*_ and *σ*_*C*max,*j*_ using equations (D.3) gives

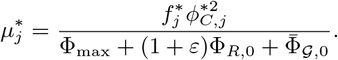

Now 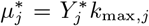 and 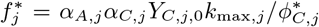, from equation (2.9), and on substituting for 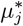 and 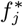 into the above and rearranging we obtain

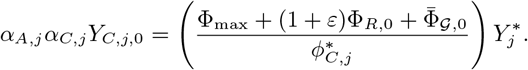

This can now be substituted into equation (2.9) to give

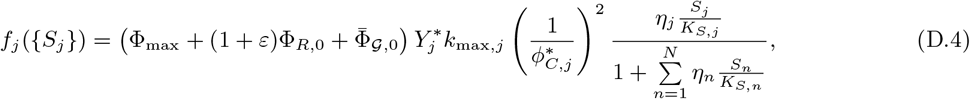

and we have eliminated the unknown constants from *f*_*j*_. In addition substituting for 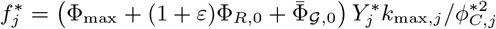 into equations (D.3) we have

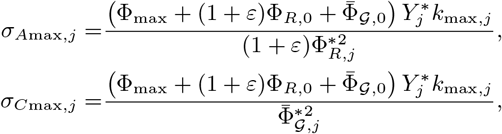

so that *σ*_*A*max,*j*_ and *σ*_*C*max,*j*_ are both expressed entirely in terms of experimentally measurable parameters.

### E Rewriting the protein dynamics equations in terms of the growth dependent protein mass fractions

The *R, A* and *C* sectors of the proteome are goverened by equations (2.19)

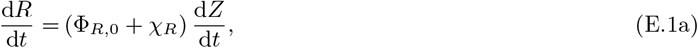

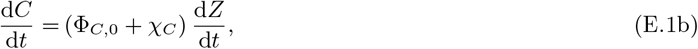

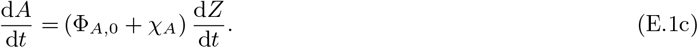

As described in section 2.2.2 we have the following relationships between total protein concentration, *Z*, and biomass concentration, *X*, and protein concentrations, *R, A* and *C*, and protein mass fractions, Φ_*R*_, Φ_*A*_ and Φ_*C*_,

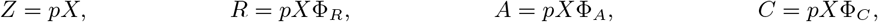

where the constant *p* is the fraction of biomass that is protein. Substituting for *R* and *Z* into equation (E.1a) we obtain

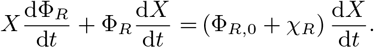

Now Φ_*R*_ = Φ_*R*,0_ + *ϕ*_*R*_, where *ϕ*_*R*_ is the growth dependent ribosomal mass fraction, and we use this expression to substitute for Φ_*R*_ into the above equation giving

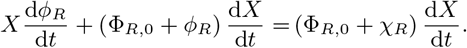

Rearranging we obtain

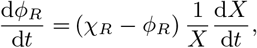

and as the growth rate, *μ* = (1*/X*)(d*X/*d*t*) we have

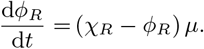

We use the same method to rewrite the equations for *C* and *A*, equations (E.1b) and (E.1c), in terms of *ϕ*_*C*_ and *ϕ*_*A*_ obtaining

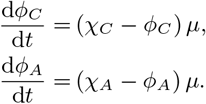

### F Governing equations and parameters used to model glucose-lactose diauxie

In the following governing equations, subscripts _1_ and _2_ denote the mass fractions and growth rates of strains *X*_1_ and *X*_2_ respectively. Subscripts _gl_ and _la_ denote parameters for growth on glucose and lactose respectively. The two strains are assumed to have the same parameters for growth on glucose. For both strains the glucose specific enzyme will always be produced so *η*_gl,1_ = *η*_gl,2_ = 1. However, the lactose specific enzyme is never produced by *X*_2_ (*η*_la,2_ = 0) and will only be produced by *X*_1_ when the concentration of glucose drops sufficiently. The point at which the lactose enzyme switches on is currently not well defined and we choose to model the switch for *X*_1_ by setting

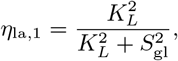

where *K*_*L*_ is a constant and *S*_gl_ is the glucose concentration. Our governing equations are

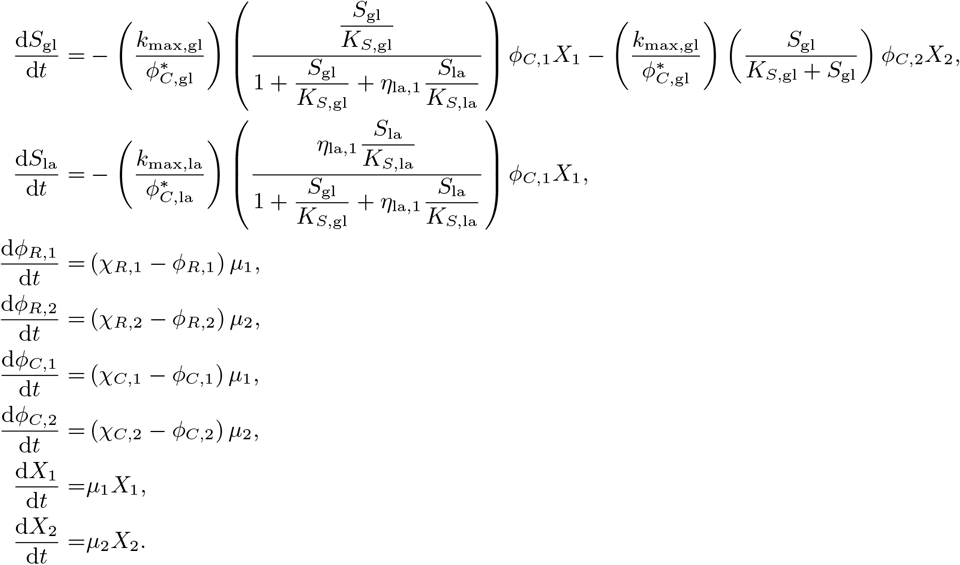

where

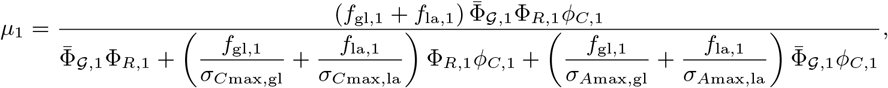

with

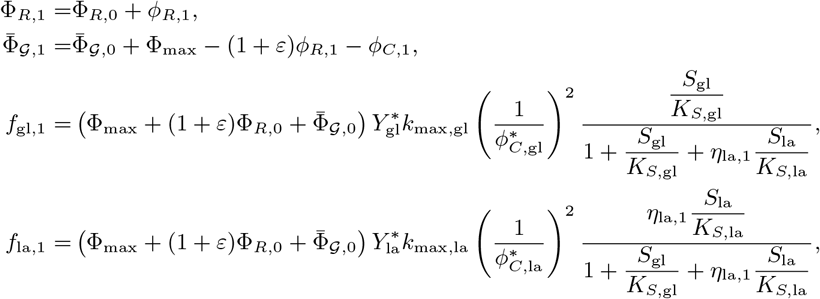

and

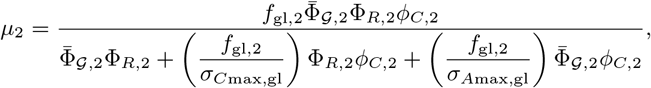

with

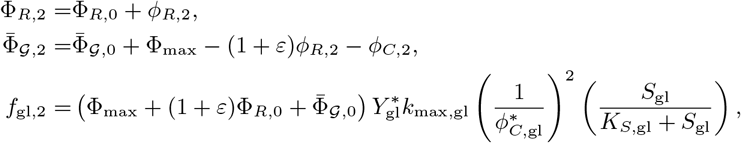

and the constants

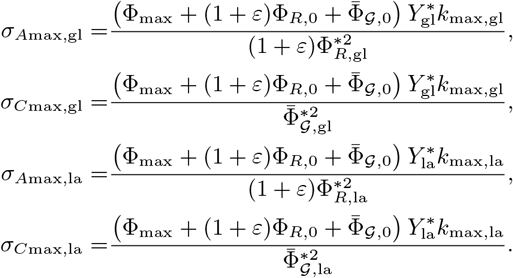

The regulation functions are given by

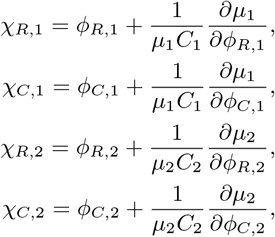

with *C*_1_ and *C*_2_ calcualated as described in Section 2.2.5.

The parameters in the governing equations whose values are taken from the literature are given in Table 4. The initial value for *ϕ*_*R*,1_ = *ϕ*_*R*,2_ was calculated from equation (2.3) with 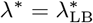. The anabolic proteins were initially assumed to be at their minimum value so *ϕ*_*A*,1_ = *ϕ*_*A*,2_ = 0 and therefore *ϕ*_*C*,1_ = Φ_max_ − (1 + *ε*)*ϕ*_*R*,1_ and similarly for *ϕ*_*C*,2_. Initial values for *S*_gl_, *S*_la_ and *X*_1_ = *X*_2_ were taken from our experimental data at *t* = 0.

**Table 4:** Parameters for *E. coli* taken from the literature

| Parameter | Description | Value |
| --- | --- | --- |
| $60 \ln(2)/\lambda_{LB}^*$ | Doubling rate on Luria-Bertani broth | 25 (mins) <sup>a</sup> |
| $\lambda_{gl}^*$ | Growth rate on glucose | 0.76 (h<sup>-1</sup>) <sup>b</sup> |
| $\lambda_{la}^*$ | Growth rate on lactose | 0.68 (h<sup>-1</sup>) <sup>b</sup> |
| $K_{S,gl}$ | Half saturation constant on glucose | 0.04 (gL<sup>-1</sup>) <sup>b</sup> |
| $K_{S,la}$ | Half saturation constant on lactose | 0.43 (gL<sup>-1</sup>) <sup>b</sup> |
| $\Phi_{R,0}$ | Minimum ribosomal mass fraction | 0.05 <sup>c</sup> |
| $\nu_R$ | Fitted parameter in growth law | 11.0 (h<sup>-1</sup>) <sup>c</sup> |
| $\lambda_L$ | Fitted parameter in growth law | 1.17 (h<sup>-1</sup>) <sup>c</sup> |
| $\Phi_{max}$ | Maximum mass fraction of growth dependent proteins | 0.43 <sup>d</sup> |
| $\varepsilon$ | Constant relating uninduced sector to R-sector | 0.3 <sup>d</sup> |

<sup>a</sup>Data from [22].<sup>b</sup>Data from [37].<sup>c</sup>Data from [18].<sup>d</sup>Data from [11].

The parameters *Y*_gl_, *Y*_la_, Φ _𝒢,0_ and *K*_*L*_ were determined by least-squares curve fitting to give a best fit to our experimental data (see Table 5). Fitting was performed using the Matlab function lsqcurvefit [28].

**Table 5:** Fitted parameters.

| Parameter | Description | Best fit value |
| --- | --- | --- |
| $Y_{gl}$ | Yield on glucose ((OD600 $X$)/(g/L $S_{gl}$)) | 1.33 |
| $Y_{la}$ | Yield on lactose ((OD600 $X$)/(g/L $S_{la}$)) | 0.80 |
| $\Phi_{G,0}$ | Minimum mass fraction of $\Phi_G$ | 10<sup>-8</sup> |
| $K_L$ | Constant in function regulating lactose uptake (g/L) | 10<sup>-4</sup> |

## Notes

### Competing Interest Statement

The authors have declared no competing interest.

### Summary of Updates

Original submission was missing references

## References

1. Monod J. Recherches sur la croissance des cultures bactériennes. PhD thesis. Paris: Hermann: Sciences naturelles: Université de Paris, 1942

2. Monod J. The growth of bacterial cultures. Annual Review of Microbiology 1949; 3:371–94. doi: 10.1146/annurev.mi.03.100149.002103

3. Chu D and Barnes DJ. The lag-phase during diauxic growth is a trade-off between fast adaptation and high growth rate. Scientific Reports 2016; 6. doi: 10.1038/srep25191

4. Kompala DS, Ramkrishna D, and Tsao GT. Cybernetic modeling of microbial growth on multiple substrates. Biotechnology and bioengineering 1984; 26:1272–81. doi: 10.1002/bit.260261103

5. Salvy P and Hatzimanikatis V. Emergence of diauxie as an optimal growth strategy under resource allocation constraints in cellular metabolism. Proceedings of the National Academy of Sciences 2021; 118. doi: 10.1073/pnas.2013836118

6. Görke B and Stülke J. Carbon catabolite repression in bacteria: many ways to make the most out of nutrients. Nature Reviews Microbiology 2008; 6:613–24. doi: 10.1038/nrmicro1932

7. Aggarwal RK and Narang A. Inducer exclusion, by itself, cannot account for the glucose-mediated lac repression of Escherichia coli. Biophysical Journal 2022; 121:820–9. doi: 10.1016/j.bpj.2022.01.016

8. Hogema BM, Arents JC, Bader R, Eijkemans K, Inada T, Aiba H, and Postma PW. Inducer exclusion by glucose 6-phosphate in Escherichia coli. Molecular microbiology 1998; 28:755–65. doi: 10.1046/j.1365-2958.1998.00833.x

9. Traxler MF, Chang DE, and Conway T. Guanosine 3’,5’-bispyrophosphate coordinates global gene expression during glucose-lactose diauxie in Escherichia coli. Proceedings of the National Academy of Sciences 2006; 103:2374–9. doi: 10.1073/pnas.0510995103

10. Scott M, Klumpp S, Mateescu EM, and Hwa T. Emergence of robust growth laws from optimal regulation of ribosome synthesis. Molecular systems biology 2014; 10:747. doi: 10.15252/msb.20145379

11. You C, Okano H, Hui S, Zhang Z, Kim M, Gunderson CW, Wang YP, Lenz P, Yan D, and Hwa T. Coordination of bacterial proteome with metabolism by cyclic AMP signalling. Nature 2013 Aug; 500:301–6. doi: 10.1038/nature12446

12. Giordano N, Mairet F, Gouzé JL, Geiselmann J, and De Jong H. Dynamical allocation of cellular resources as an optimal control problem: novel insights into microbial growth strategies. PLoS computational biology 2016; 12:e1004802. doi: 10.1371/journal.pcbi.1004802

13. Ibarra RU, Edwards JS, and Palsson BO. Escherichia coli K-12 undergoes adaptive evolution to achieve in silico predicted optimal growth. Nature 2002; 420:186–9. doi: 10.1038/nature01149

14. Scott M, Gunderson CW, Mateescu EM, Zhang Z, and Hwa T. Interdependence of Cell Growth and Gene Expression: Origins and Consequences. Science 2010; 330:1099–102. doi: 10.1126/science.1192588

15. Maitra A and Dill KA. Bacterial growth laws reflect the evolutionary importance of energy efficiency. Proceedings of the National Academy of Sciences 2015; 112:406–11. doi: 10.1073/pnas.1421138111

16. Weiße AY, Oyarzún DA, Danos V, and Swain PS. Mechanistic links between cellular trade-offs, gene expression, and growth. Proceedings of the National Academy of Sciences 2015; 112:E1038–E1047. doi: 10.1073/pnas.1416533112

17. Pavlov MY and Ehrenberg M. Optimal control of gene expression for fast proteome adaptation to environmental change. Proceedings of the National Academy of Sciences 2013; 110:20527–32. doi: 10.1073/pnas.1309356110

18. Erickson D, Schink SJ, Patsalo V, Williamson JR, Gerland U, and Hwa T. A global resource allocation strategy governs growth transition kinetics of Escherichia coli. Nature 2017 Oct; 551:119–23. doi: 10.1038/nature24299

19. Basan M, Honda T, Christodoulou D, Hörl M, Chang YF, Leoncini E, Mukherjee A, Okano H, Taylor BR, Silverman JM, et al. A universal trade-off between growth and lag in fluctuating environments. Nature 2020; 584:470–4. doi: 10.1038/s41586-020-2505-4

20. Kremling A, Geiselmann J, Ropers D, and Jong H de. An ensemble of mathematical models showing diauxic growth behaviour. BMC systems biology 2018; 12:1–16. doi: 10.1186/s12918-018-0604-8

21. Swinnen I, Bernaerts K, Dens E, Geeraerd A, and Van Impe J. Predictive modelling of the microbial lag phase: a review. International Journal of Food Microbiology 2004; 94:137–59. doi: 10.1016/j.ijfoodmicro.2004.01.006

22. Tao H, Bausch C, Richmond C, Blattner FR, and Conway T. Functional genomics: expression analysis of Escherichia coli growing on minimal and rich media. Journal of bacteriology 1999; 181:6425–40. doi: 10.1128/JB.181.20.6425-6440.1999

23. Scott M and Hwa T. Bacterial growth laws and their applications. Current Opinion in Biotechnology 2011; 22:559–65. doi: 10.1016/j.copbio.2011.04.014

24. Hui S, Silverman JM, Chen SS, Erickson DW, Basan M, Wang J, Hwa T, and Williamson JR. Quantitative proteomic analysis reveals a simple strategy of global resource allocation in bacteria. Molecular Systems Biology 2015; 11:784. doi: 10.15252/msb.20145697

25. Mostovenko E, Deelder A, and Palmblad M. Protein expression dynamics during Escherichia Coli glucose-lactose diauxie. BMC Microbiology 2011; 11. doi: 10.1186/1471-2180-11-126

26. Brown T. Gene Cloning and DNA Analysis, An Introduction. 6th edition. English. 6th. United Kingdom: John Wiley & Sons Ltd, 2010

27. Motulsky H. GraphPad Software. https://www.graphpad.com. San Diego, California USA, 2021

28. MATLAB. MATLAB version 9.8.0.1359463 (R2020a) Update 1. The Mathworks, Inc. Natick, Massachusetts, 2020

29. Jaishankar J and Srivastava P. Molecular basis of stationary phase survival and applications. Frontiers in microbiology 2017; 8:2000. doi: 10.3389/fmicb.2017.02000

30. Spencer CC, Bertrand M, Travisano M, and Doebeli M. Adaptive diversification in genes that regulate resource use in Escherichia coli. PLoS genetics 2007; 3:e15. doi: 10.1371/journal.pgen.0030015

31. Wang J, Atolia E, Hua B, Savir Y, Escalante-Chong R, and Springer M. Natural variation in preparation for nutrient depletion reveals a cost–benefit tradeoff. PLoS biology 2015; 13:e1002041. doi: 10.1371/journal.pbio.1002041

32. New AM, Cerulus B, Govers SK, Perez-Samper G, Zhu B, Boogmans S, Xavier JB, and Verstrepen KJ. Different levels of catabolite repression optimize growth in stable and variable environments. PLoS biology 2014; 12:e1001764. doi: 10.1371/journal.pbio.1001764

33. Siegal ML. Shifting sugars and shifting paradigms. PLoS biology 2015; 13:e1002068. doi: 10.1371/journal.pbio.1002068

34. Mori M, Marinari E, and De Martino A. A yield-cost tradeoff governs Escherichia coli’s decision between fermentation and respiration in carbon-limited growth. NPJ systems biology and applications 2019; 5:1–9. doi: 10.1038/s41540-019-0093-4

35. Wang X, Xia K, Yang X, and Tang C. Growth strategy of microbes on mixed carbon sources. Nature communications 2019; 10:1–7. doi: 10.1038/s41467-019-09261-3

36. Murray JD. Mathematical Biology. Springer Berlin Heidelberg, 2013

37. Doshi P and Venkatesh K. An optimal model for microbial growth in a multiple substrate environment: simultaneous and sequential utilization. Process Biochemistry 1998; 33:663–70. doi: 10.1016/S0032-9592(98)00031-4

